# Host prior exposure augments heterogeneity in gene expression in both host and pathogen during *in vivo* infection

**DOI:** 10.1101/2025.08.20.671316

**Authors:** Anna A. Pérez-Umphrey, Jeremy Miller, Edan R. Tulman, Jesse Garrett-Larsen, Michal Vinkler, Kate Langwig, Steven J. Geary, James S. Adelman, Dana M. Hawley

## Abstract

Variability in acquired protection, whether from prior pathogen exposure or vaccination, is increasingly recognized as a key determinant of host population-level variation in disease traits. It remains unclear whether this extends to the within-host physiological environment and what the consequences are for reinfecting pathogens. Here, we asked whether prior pathogen exposure of hosts induces gene expression heterogeneity in the host and/or pathogen during infection. We quantified gene expression *in vivo* following high-dose pathogen challenge of house finches (*Haemorhous mexicanus*) previously given controlled, varied exposure histories to a bacterial pathogen (*Mycoplasma gallisepticum*; MG). To measure gene expression heterogeneity, we collected transcriptomic data from two host tissues (conjunctiva and spleen), and, simultaneously, from pathogen infecting the primary site of infection (conjunctiva). In the conjunctiva, but not the spleen, prior pathogen exposure induced significant heterogeneity in host gene expression relative to pathogen-naïve hosts. Further, hosts that received a lower prior exposure dose rather than a higher primary dose showed the greatest within-group heterogeneity in expression during re-challenge. Functional enrichment analyses for significantly variable host genes indicated an over-representation of terms involved in the immune system’s response to pathogens, namely a diversified inflammatory response, in birds with prior pathogen exposure. The infecting pathogen from the conjunctiva followed similar patterns of heterogeneity in host gene expression, where pathogen infecting hosts with prior exposure had more heterogeneous expression than those infecting pathogen-naïve hosts. While the exact mechanisms that underlie greater variation in gene expression cannot be resolved by this study, our results are consistent with the hypothesis that prior host exposure induces a within-host environment that promotes heterogeneous gene expression across both hosts and pathogens. This suggests that to understand the coevolutionary dynamics of infectious diseases we must consider not only the genetic sequence variation, but also gene expression variation in host and pathogen.

## Introduction

A primary focus of disease and evolutionary ecology is to characterize the reciprocal selective pressures of host-pathogen interactions (Ebert & Bull, 2008), and the processes by which this antagonistic coevolutionary relationship generates host and pathogen diversity. In nature, host populations are seldom homogenous in their defenses (Ebert & Bull, 2008), and this heterogeneity can have significant epidemiological (Elie et al., 2022; Gomes et al., 2022; Lloyd et al., 2020; Vazquez-Prokopec et al., 2016) and evolutionary (Ebert & Bull, 2008; Fleming-Davies et al., 2015; P. S. White et al., 2020) consequences for host-pathogen dynamics. Because genetic differences among hosts (Aalen et al., 2015; Gibson & Lively, 2019; L. A. White et al., 2020) are a key source of inter-individual disease trait variation (Lazzaro & Little, 2009), substantial research has focused on the mitigating or amplifying effects of host genetic diversity on pathogen prevalence and adaptation to host populations (Ganz & Ebert, 2010; Gibson & Lively, 2019; Morley et al., 2017; P. S. White et al., 2020). However, the extent to which host gene regulatory variation influences gene expression of the infecting pathogen, or vice versa, remains largely unexplored.

The within-host environment serves as the “pathogen’s habitat” and is a key source of selective pressures (Cobey, 2014; Osnas & Dobson, 2012). The ability of a pathogen to effectively replicate and evade immune responses within the host can result in trade-offs for within- and between-host processes such as pathogenicity and transmission, all of which must be considered to understand a pathogen’s evolutionary trajectory (Osnas & Dobson, 2012). Host genetic heterogeneity can act as a cost to components of pathogen fitness both within- and between-hosts. For example, host population genetic diversity, whether genome-wide or at immune-specific loci, is associated with pathogen resistance in several systems (e.g., Savage and Zamudio 2011; Budischak et al., 2023), and genetic diversity can mitigate the ability of pathogens to transmit among hosts (i.e., the “monoculture effect”; Ekroth et al., 2019). This limits pathogen prevalence (Ganz & Ebert, 2010), local adaptation or specialization to hosts (Morley et al., 2017; P. S. White et al., 2020), and evolution of virulence, even where there is a strong selective advantage for it (P. S. White et al., 2020). Although population-level host genetic heterogeneity can limit pathogen prevalence and potential for local adaptation, it can simultaneously augment pathogen strain or genetic diversity due to the variable selective pressures that heterogeneous hosts place on the pathogen. For example, the coexistence of multiple malaria strains is facilitated by different host genotypes that favor the within-host competition of one strain at the expense of another (de Roode et al., 2004). In terms of pathogen genetic diversity, bacterial lineages infecting nematodes accumulated significantly more whole-genome and per-gene diversity when passaged through hosts with variable genetic defenses, relative to homogenous nematode populations (Hoang et al., 2024). Likewise, pathogen diversity determines the protective effectiveness of host population genetic heterogeneity (Ganz & Ebert, 2010; van Baalen & Beekman, 2006), and can maintain host genetic variation at immune loci (Lazzaro & Little, 2009).

Heterogeneity in a host population’s disease traits can also be environmentally induced (Ben-Ami et al., 2010; Dwyer et al., 1997; Gomes et al., 2014) and, as such, gene expression profiles may contribute to patterns of reciprocal selective pressures between hosts and pathogens (Barribeau et al., 2014). This is highly salient given that all infections occur within an environmental context, to which host loci must be able to respond somewhat plastically if they are to optimize immunological costs/benefits (Råberg et al., 1998; Schmid-Hempel, 2003). Such heterogeneity in gene expression during active infection, whether for immune genes or other functions relevant to a pathogen’s response, may ultimately create a more diverse within-host selective environment for pathogens.

Pathogen exposure history is increasingly recognized as an important ecological variable that can augment host population-level heterogeneity in traits relevant to transmission dynamics, such as susceptibility (Dwyer et al., 1997; Gomes et al., 2014; Hawley et al., 2024; Langwig et al., 2017). In nature, pathogen exposure dose and frequency likely vary (Aiello et al., 2016; Leon & Hawley, 2017; Turner et al., 2016), however, even when exposure dose is controlled (e.g., vaccination), the degree and longevity of the resulting immune protection can vary widely among individuals (Le et al., 2021), such that the host population a pathogen encounters has heterogenous standing protection (Hawley et al., 2024; Langwig et al., 2017; Weitzman et al., 2022). How these disease traits manifest is inextricably linked to the within-host environment and the pathogen’s success within the host (Mideo et al., 2008); however, studies of host heterogeneity rarely consider parasite variability in tandem (Ganz & Ebert, 2010).

The house finch (*Haemorhous mexicanus*) and its endemic bacterial pathogen (*Mycoplasma gallisepticum*) is a relevant disease system to address these questions given that host acquired immune protection is incomplete even after high initial exposure doses (Fleming-Davies et al., 2018; Weitzman et al., 2022), such that reinfections are common (Leon & Hawley, 2017). Originally a respiratory pathogen of poultry, MG’s emergence in house finches in the early 1990s resulted in significant population declines (Dhondt et al., 1998). Infection in house finches causes severe inflammation of the conjunctiva (Ley et al., 1996), which can be indirectly fatal by making the host more susceptible to predation (Adelman et al., 2017). Recent work in this system has shown that prior exposure to *Mycoplasma gallisepticum* (MG) augments heterogeneity in disease traits relevant to between-host dynamics upon reinfection, such as susceptibility (Hawley et al., 2024; Pérez-Umphrey et al., 2025 [Preprint]) and pathogen loads (Garrett-Larsen et al., 2025 [Preprint]), particularly at low doses (Leon et al., 2019). However, it remains unknown if prior exposure to MG in house finches also augments heterogeneity in gene regulatory responses, and whether pathogen gene expression covaries. This might be due to either the direct influence of prior exposure treatment on the pathogen, or, indirectly, variation in the within-host environment and selective pressures in the form of variable host standing immunity (Cobey, 2014).

Here, we quantified host and pathogen gene expression *in vivo*, during high-dose challenges of house finches given variable MG exposure histories (no prior MG, a prior low-dose of MG, or prior intermediate-dose of MG). We collected transcriptomic datasets for two host lymphoid tissues (conjunctiva and spleen) and for infecting MG isolated from the primary site of infection (the conjunctiva), to ask whether prior pathogen exposure increases heterogeneity of the within-host environment and if pathogen expression covaries during reinfection.

## Methods

### Experimental design

Wild-caught (but MG-naive) house finches (n = 27) were randomly assigned to one of six treatment groups in a 3 x 2 factorial experimental design, where sex ratios were kept as even as possible across treatments (Table 1). Variation in host prior exposure history was created by inoculating birds with different initial pathogen doses (sham control [no MG], low, or intermediate MG dose). The birds were allowed to recover and were then either reinfected with a high dose of MG or given a sham control (Figure 1). All inoculations were done with the index isolate of *Mycoplasma gallisepticum* (VA1994; Ley et al., 1996). Three days post-secondary challenge, two host tissues – a periocular lymphoid tissue that is the primary site of infection (palpebral conjunctiva) and a remote secondary lymphoid tissue (spleen) – were collected from the host to generate Poly(A)-RNA-sequencing libraries. For birds in the high secondary challenge treatment only, the conjunctival tissue was also used to extract the infecting MG to generate total RNA-sequencing libraries.

**Figure 1.**
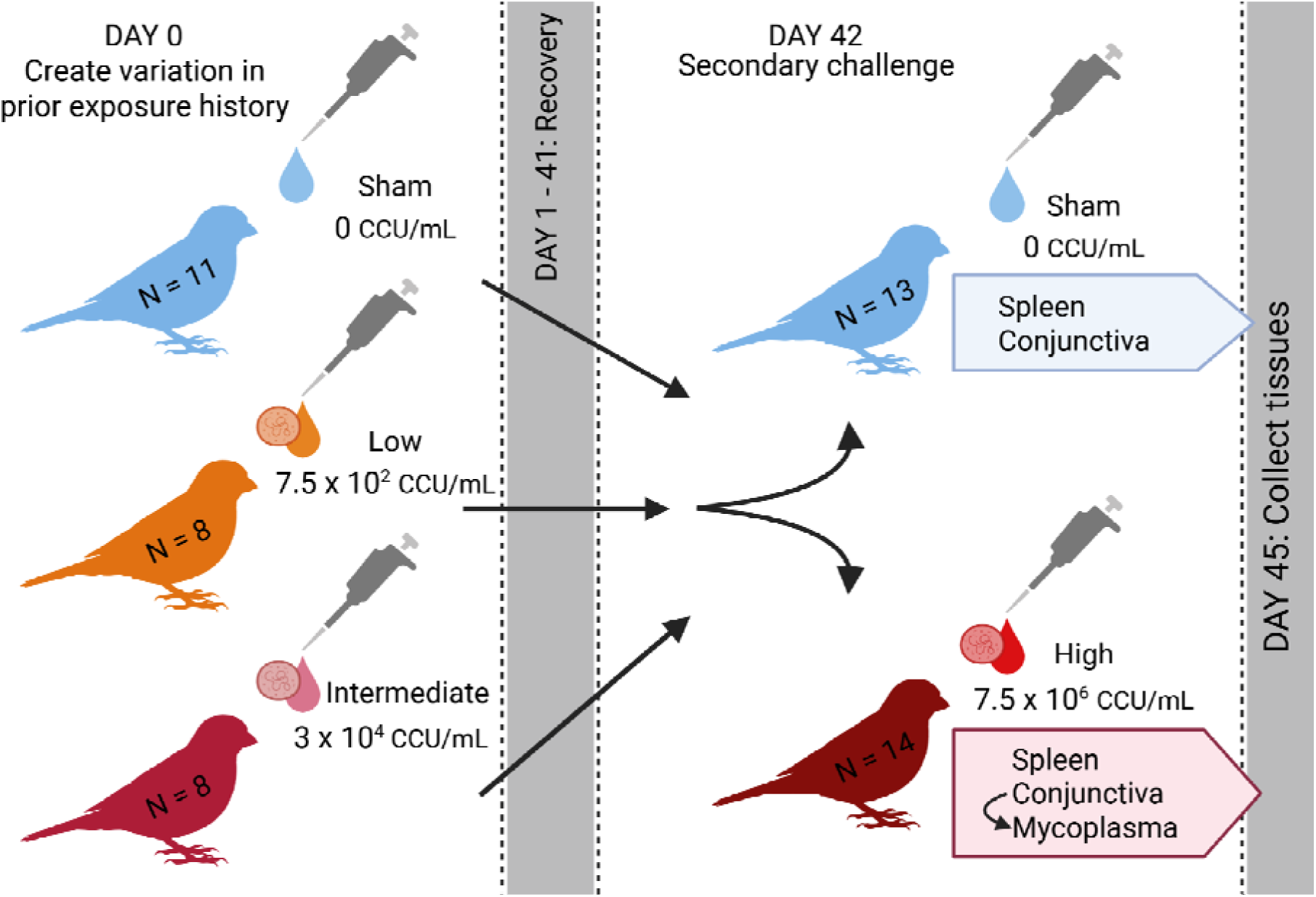
Experimental design. Birds received one of three initial exposure doses (sham [no MG], low, or intermediate) of the same MG strain via ocular inoculation. Birds were allowed to recover for forty-one days and then were re-challenged with one of two secondary doses (sham or high dose). Samples were collected three days post-secondary challenge and used to generate RNA-seq libraries. Spleen and conjunctiva tissue were collected for each bird and pathogen (“*Mycoplasma*” above) was sequenced only from the conjunctiva of birds that received a high secondary dose. Created in BioRender.

**Table 1.** Treatment groups and sample sizes. Birds were randomly placed in one of six treatment groups, which were a combination of three primary (sham, low, intermediate) and two secondary treatments (sham, high).

| Inoculation Treatments |  | Sex per treatment |  |  |
| --- | --- | --- | --- | --- |
| Primary | Secondary | Female | Male | Total |
| Sham | Sham | 2 | 3 | 5 |
| Sham | High | 3 | 3 | 6 |
| Low | Sham | 2 | 2 | 4 |
| Low | High | 2 | 2 | 4 |
| Int | Sham | 2 | 2 | 4 |
| Int | High | 2 | 2 | 4 |
| TOTAL |  | 13 | 14 | 27 |

### House finch capture, quarantine, and housing

Birds were captured on Virginia Tech’s campus and nearby residential areas of Blacksburg, Virginia, USA in the summer of 2022 using mist-nets and wire-cage traps at baited feeders. Only hatch-year birds (identified by plumage coloration and yellow gapes) were retained as they have the lowest chance of MG exposure in the wild. Upon capture, birds received a captive metal band and sex was determined by plumage coloration after birds had molted (late-August/early-September). During a twenty-one-day quarantine, birds were pair-housed in 46 x 76 x 46 (cm) cages in animal care facilities on Virginia Tech’s campus. During this time they were periodically assessed for the development of any clinical signs of MG infection (described below) that can result from potential exposure just prior to capture. In the final week of quarantine, birds had blood drawn from the brachial vein and their sera tested for antibodies indicating previous MG exposure. These screenings were performed via *M. gallisepticum* antibody enzyme-linked immunosorbent assays (IDEXX, Westbrook, ME) using an optical density (OD450) threshold of 0.061 (Hawley et al., 2011). Birds were not used in the study if they were seropositive, developed clinical signs of disease during quarantine, or had a cagemate that was at any point clinically diseased. Selected animals were moved into clean cages and single-housed for the duration of the study.

### Inoculations and sample collection

Birds were inoculated (day 0 of experiment - timeline in Table S1) with one of three primary MG doses (sham control [0 CCU/mL], low [7.5 × 10^2^ CCU/mL], or intermediate [3 × 10^4^ CCU/mL]). Inocula were prepared from isolates of VA1994 (Ley et al., 1996) and diluted in sterile Frey’s media to reach desired concentrations, then delivered via ocular absorption using micropipettes to dispense 35 µl per eye of the requisite dose. Birds were allowed to recover for forty-one days between their primary and secondary challenges. This interval was sufficient for birds to recover from initial infection, which was confirmed by the complete resolution of pathology and a low or absent pathogen load. On day 42 post-primary inoculation (DPPI 42) birds in each primary treatment group received one of two possible secondary treatments: a sham control (sterile Frey’s media, 0 CCU/mL) or a high dose (7.5 × 10^6^ CCU/mL) of the same MG strain used in the primary challenge. Clinical signs (eyescore) and pathogen load (eyeswab) data were collected longitudinally for each bird (Table S1). Because conjunctivitis presents as highly visible swelling, crusting, and exudate, it can be categorically scored in terms of severity from 0-3 per eye for a total “eyescore” (left plus right eye) of 0-6 per a given bird (Sydenstricker et al., 2005). The same experienced observer (co-author DMH) blind to treatment group collected these data throughout the experiment. Pathogen load data were collected by swabbing palpebral conjunctiva with sterile cotton-tipped swabs which were then swirled in 300 µl tryptose phosphate broth and discarded. This was done for each eye using separate swabs pooled in the same sample tube. MG DNA was extracted from this solution and quantified via qPCR assay (see Supplemental Methods).

Three days post-secondary inoculation, eyeswabs and eyescore data were collected, after which birds were euthanized by exposure to isoflurane followed by cervical dislocation and decapitation. Dissections and tissue collection were performed immediately post-mortem. Tools and workspaces were decontaminated with 70% ethanol and RNaseZAP™ (Invitrogen, Carlsbad, CA) between processing each bird. Each eye’s lower palpebral conjunctiva (stored together in one tube) and spleen were excised and placed in sterile cryovials containing 1 mL of TRIzol™ reagent (Invitrogen, Carlsbad, CA). Spleens were frozen on dry ice and later moved to -80°C. To “wash” conjunctival tissue and suspend MG into solution, conjunctiva were vortexed for 10-15s and stored at 4°C for 72 hours. The samples were then moved to -80°C for storage until nucleic acid extraction.

### Library preparation and sequencing

Library preparations and sequencing runs were performed at the Center for Genome Innovation, Institute for Systems Genomics, University of Connecticut. Raw sequencing data is available on NCBI’s Single Read Archive (SRA); BioProject accession: PRJNA1298300.

#### Host

Conjunctiva and spleen tissue were thawed and manually homogenized with sterile, RNase-free pellet pestles for ∼1 minute while in TRIzol™ solution. Total RNA was extracted using a Zymo Direct-zol Miniprep Kit (Zymo Research Corporation, Irvine, CA) during which samples were treated with DNase I, per manufacturer recommendations. Total RNA sample quality was checked by TapeStation High Sensitivity RNA ScreenTape Analysis prior to sequencing (minimum input of 200 ng and ≥7 RIN). Since mycoplasmas lack poly-adenylated RNAs (Portnoy & Schuster, 2008), isolating eukaryotic poly-adenylated mRNAs during library construction effectively prevents MG RNA contamination in host libraries. Libraries were prepared using Illumina Stranded mRNA Prep Ligation kits (Illumina, San Diego, CA) and were sequenced with 20M 150 bp paired-end reads on an Illumina NovaSeq 6000 (S4 300 cycle version 1.5 sequencing kit). Conjunctiva and spleen tissue were sequenced on two separate sequencing runs and analyzed separately (below) to avoid any potential sequencing batch effects. ***Pathogen***

Non-poly-adenylated *Mycoplasma gallisepticum* RNA was enriched by negatively selecting out, first, host mRNA in a library pre-step (NEBNext® Poly(A) mRNA Magnetic Isolation Module, NEB, Ipswich, MA). Then, host and bacterial rRNA were removed during sequencing library preparation using the Illumina Stranded Total RNA Prep, Ligation with Ribo-Zero™ Plus kit (Illumina, San Diego, CA). These MG-enriched libraries were sequenced on an Illumina NovaSeq 6000 (S4 300 cycle version 1.5 sequencing kit) targeting 7M 150 bp paired-end reads per sample.

### Data analysis

#### Bioinformatic processing

Data were demultiplexed by the sequencing facility and all bioinformatic steps were performed on Virginia Tech’s Advanced Research Computing TinkerCliffs cluster. All sequencing data (with adjustments made for host versus pathogen analyses – described below) were processed via a snakemake pipeline “RNASeq-snakemake-pipeline” (https://github.com/JoannaGriffiths) designed to automate computational steps from raw sequence data input to read count quantification file generation, whose steps are briefly described here: Sequence read quality control and adapter removal was performed for each forward and reverse FASTQ file using FastQC v. 0.11.9 (https://www.bioinformatics.babraham.ac.uk/projects/fastqc/) and fastp v. 0.23.2 (Chen et al., 2018). Reads were filtered according to default fastp v. 0.23.2 parameters, which includes polyG tail trimming for NovaSeq-generated data. The house finch annotated reference genome (“HaeMex1”) and transcriptome were accessed via NCBI (accession: GCF_027477595.1; https://www.ncbi.nlm.nih.gov/datasets/genome). The transcriptome was indexed (decoy-aware) in Salmon v. 1.9.0 (Patro et al., 2017). Then, filtered reads were aligned to the transcriptome and quantified in “mapping-based mode” in Salmon v. 1.9.0 (Patro et al., 2017) with the optional --*validatemappings*, --*gcBias*, and --*seqBias* commands. The genome of the *Mycoplasma gallisepticum* isolate (“VA94_7994-1-7P”) used in this study was likewise accessed through NCBI (accession: GCF_000286675.1; https://www.ncbi.nlm.nih.gov/datasets/genome). Its transcriptome was extracted from the accessed GFF file using GffRead v. 0.12.7 (Pertea & Pertea, 2020) and then indexed in Salmon (decoy-unaware). Filtered reads were aligned and quantified in mapping-based mode in Salmon v. 1.9.0 with the --v*alidatemappings* option.

Once read count files were generated, they were imported into R (v. 4.3.1; R Core Team 2023) with *tximport* 1.30.0 (Soneson et al., 2016). Downstream analyses were performed using Bioconductor software package *DESeq2* v. 1.42.0 (Love et al., 2014). Transcript and gene metadata were incorporated by creating a TxDb object with accessed GTF files from NCBI and using the ‘makeTxDbFromGFF’ function in *GenomicFeatures* v. 1.54.4 (Lawrence et al., 2013). Genes with fewer than ten reads across all samples were removed from each dataset (conjunctiva: 2,234 removed and 17,032 retained; spleen: 2,665 removed and 16,601 retained; *Mycoplasma*: 64 removed and 647 retained) and data were r-logged (described below) and normalized by sample-specific size factors (Love et al., 2014). The DESeqDataset design formula was ∼Treatment, where treatment was one of the six possible combinations of primary (sham, low, intermediate) and secondary challenge (sham or high). However, MG sample sequencing depth depended on the pathogen load of each bird which is inherently confounded with its treatment group (Figure S1). To account for this, we created surrogate variables using the *sva* v. 3.50.0 package (Leek 2014) and then controlled for those variables by adding them to the DESeq model for the MG dataset (see Supplementary-Results for numbers of assigned reads).

Raw RNA-seq count data typically show greater variances for genes with higher mean expression. While a log_2_-scale-transformation can account for this, it may then inflate the variance of genes with low mean expression (Love et al., 2014). To ensure our data were homoscedastic, we used the regularized-logarithmic (rlog) transformation available in DESeq2. This transforms the counts (already normalized by library size) to a log_2_ scale and employs a Bayesian “shrinkage” procedure to model the data: it assigns a unique term to each sample and incorporates a prior distribution on the coefficients, which is inferred from the data (Love et al., 2014). This stabilizes variances and removes their dependence on the mean. These transformed data were used for all downstream analyses.

#### Tests of heterogeneity

To ask whether prior pathogen exposure induced heterogeneity in gene expression upon secondary challenge, we first removed genes with little to no variance in expression across all samples. This is a common filtering step (van Iterson et al., 2010; Zehetmayer et al., 2022) used to exclude low-information genes with non-experimentally relevant expression. For each tissue, we retained those genes that had a variance greater than 0.5 (Figure S2; n = 462 conjunctival genes; n = 177 splenic genes; n = 413 MG genes), rather than a pre-specified arbitrary number of genes, to facilitate the comparability of heterogeneity in gene expression across sample types. The analysis was also performed using ±10% the 0.5 variance threshold. This was done to ensure that our selection of variance threshold did not inordinately influence the downstream analyses and overall conclusions [See Supplementary Results - Tables S2 and S3].

With these filtered datasets, we then performed a permutational multivariate analysis of variance (PERMANOVA; n = 1,000 permutations) of the r-logged data. PERMANOVA is a resemblance-based statistical test for non-parametric multivariate data which tests the null hypothesis that group centroids are equivalent within a given resemblance measure space (Anderson, 2001; Anderson & Walsh, 2013). This was implemented using the ‘adonis2’ function in *vegan* v. 2.6.4 (Oksanen et al., 2022) and specifying a Manhattan distance matrix. A Manhattan distance matrix, which uses absolute distances, is appropriate because it resolves two common attributes of RNA-seq datasets: it preserves discriminative power as data dimensionality increases (Aggarwal et al., 2001) and is robust to zero-inflation (Skinnider et al., 2019). This was followed by a test for multivariate dispersion homogeneity with n = 1,000 permutations (PERMDISP; Anderson 2006) using the ‘betadisper’ function in *vegan* v. 2.6.5 (Oksanen et al., 2022). Post-hoc comparisons were performed with Tukey HSD tests.

#### Functional enrichment analyses

To identify those loci in both host and pathogen that are differentially heterogeneous according to host exposure history, we used the Brown-Forsythe test to compare variances per gene (Huang et al., 2020), using the same filtered, r-logged datasets used to test for differences in within-group homogeneity (above). The Brown-Forsythe is a modification of Levene’s test of homogeneity and uses the median rather than the mean to test for equality in group variances, and is therefore more robust to data non-normality and outliers (Brown & Forsythe, 1974). This analysis was performed for conjunctival tissue and the infecting MG. Analyses compared gene variances between groups of birds that received the same secondary challenge (high dose) but were either naïve (sham+high) or had prior pathogen exposure (low+high, intermediate+high) at the time of the challenge. P-values were adjusted using a false discovery rate (FDR; Benjamini & Hochberg 1995) of 0.1 (we report results at a 0.05 cutoff as well) to identify genes that were significantly differentially variable (SDV) between groups. These were used to create gene lists for downstream functional enrichment analyses. The more lenient FDR of 0.1 was used at this step to generate larger gene lists so that relevant biological pathways would not be missed. This is a relatively common approach [e.g. (Bonisoli-Alquati et al., 2020; Kumar et al., 2021; Tandoh et al., 2022; Zhai et al., 2014)] for hypothesis generating methods, such as RNA-seq, where potentially overly-conservative cutoffs can eliminate important, novel associations (Yang et al., 2011).

A custom organism package database was created using the house finch data available on NCBI (accession: GCF_027477595.1) with *AnnotationForge* v. 1.44.0 (Carlson and Pagès 2023). Over-representation analyses for gene ontology (GO) terms for Biological Process, Molecular Function, and Cellular Component were performed in *clusterProfiler* (Wu et al., 2021), specifying an FDR (Benjamini & Hochberg, 1995) of 0.05. Enrichment analyses considered lists of genes that were identified in the Brown-Forsythe test as being SDV between groups of birds that received a high dose secondary challenge. For SDV but unannotated loci, sequence data were extracted from the GTF-converted BED file using BEDTools (Quinlan & Hall, 2010). We then used NCBI’s Nucleotide BLAST database (Basic Local Alignment Search Tool; https://blast.ncbi.nlm.nih.gov/Blast.cgi) to identify potential alignments for each sequence. Reported alignments (Table S4) are the top alignments based on E-score, percent identity, and query coverage.

For the MG dataset, we created a custom GO annotation file by extracting all predicted RefSeq Protein IDs from the GTF file and querying these for GO terms in the UniProtKB database (https://www.uniprot.org/). We then used *clusterProfiler* (Wu et al., 2021) in the same manner as that used for the house finch dataset, to test for the enrichment of GO terms belonging to those genes identified as SDV between treatment groups.

#### Pathogen loads

Differences in pathogen load by treatment group were assessed for eyeswab samples collected three days post-secondary inoculation (DPSI 3). We performed a Kruskal-Wallis test for pathogen loads (log_10_+1) of birds that received a secondary MG challenge. Eyeswabs collected from birds that received a sham secondary dose were not included in the analysis. Regardless of omnibus test significance, planned comparison Wilcoxon rank sum tests were performed to test *a priori* hypotheses about pathogen loads in treated groups relative to the sham group. Additionally, the coefficient of variation (σ / μ) was calculated per secondary MG challenge treatment group.

## Results

### Gene expression heterogeneity

#### Host

At the primary site of infection (i.e., conjunctiva), prior pathogen exposure induced significant heterogeneity in host gene expression during secondary MG challenge (PERMDISP permutation test, F = 6.09, p < 0.001; Figure 2A). Pathogen-naïve birds (sham prior exposure) were relatively uniform (mean dispersion = 226.9 [range: 191.62–293.37]) in their gene expression response to the same secondary pathogen dose and significantly differed from those that had a low prior exposure dose in a Tukey post-hoc test (sham+high vs. low+high: mean difference = -145.76, 95% CI [-241.31, -50.22], p = 0.001). The low prior dose produced the greatest within-group heterogeneity in host gene expression during secondary challenge (mean dispersion = 372.66 [range: 343.41–403.22]). The intermediate prior dose also increased heterogeneity of expression (mean dispersion = 300.88 [range: 245.54 - 445.26]), although to a lesser extent than the low+high group, and was not significantly different from the sham+high group (sham+high vs. intermediate+high: mean difference = -73.99, 95% CI [-169.53, 21.56], p = 0.193) or that of any other group (Figure 2A). Tests of dispersion showed no treatment effect (F = 0.784, p = 0.588) among spleens collected from hosts with variable exposure histories (Figure 2B).

**Figure 2.**
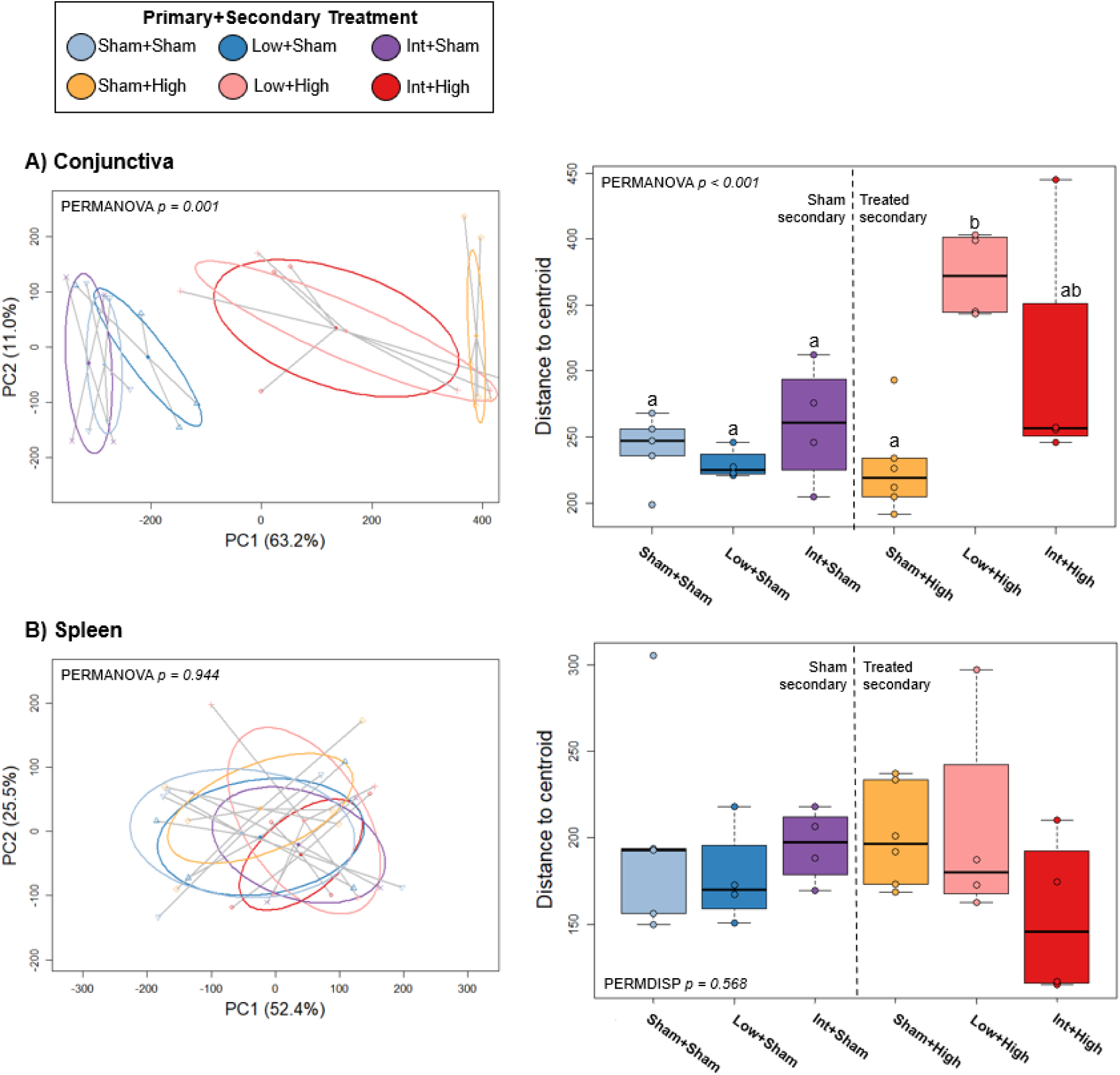
Heterogeneity in gene expression across treatments (primary + secondary) in host conjunctiva (A) and spleen (B) three days post-secondary challenge. (Left panels) PCoA plots (principal coordinate analysis) of PERMANOVA (permutational analysis of variance) results using Manhattan distance matrices of r-log-transformed gene expression data. (Right panels) Boxplots of group dispersion values shown with PERMDISP (test for multivariate dispersion homogeneity) results. Where applicable, significant post-hoc differences are represented by unshared letters between groups. The vertical dotted line highlights the data collected from hosts that received a sham versus a MG secondary dose.

In the conjunctiva, there was substantial discrimination (i.e., differing group centroids) between birds that received a secondary pathogen challenge versus those that were given a sham secondary dose (PERMANOVA permutation test, F = 6.17, p < 0.001; Figure 2A). However, groups receiving only a sham dose challenge did not differ from each other in gene expression heterogeneity regardless of their prior exposure treatment (sham+sham vs. low+sham: 11.79, 95% CI [-87.50, 111.09], p = 0.999; sham+sham vs. intermediate+sham: -18.48, 95% CI [- 117.77, 80.82], p = 0.991). In the spleen, no differences in group centroids were detected (PERMANOVA permutation test, F = 0.799 p = 0.568; Figure 2B).

#### Pathogen

Infecting pathogen from the conjunctival tissue (only collected from hosts given a MG-dose (“high”) secondary challenge) broadly followed the patterns of gene expression heterogeneity evident in its host tissue (Figure 3). Group centroids differed significantly among the three secondary-MG challenge treatments (PERMANOVA permutation test, F = 3.04, p = 0.034) and dispersion values indicated significant differences augmented by host prior pathogen exposure (PERMDISP permutation test, F = 5.58, p < 0.001). Pathogen extracted from birds that were MG-naïve at the time of challenge were more homogenous in expression response (mean dispersion = 191.95 [range: 167.03-223.13) relative to pathogen from hosts with prior low (mean dispersion = 408.12 [range: 379.00 - 440.19]) or intermediate-dose exposure (mean dispersion = 414.57 [range: 205.41-734.59]). These differences were significant in Tukey post-hoc tests, where pathogen from birds with any degree of prior exposure (low or intermediate) were more heterogeneous than that from pathogen-naïve birds (sham+high vs. low+high: -216.17, 95% CI [-428.14, -4.20], p = 0.0456; sham+high vs. intermediate+high: -222.62, 95% CI [-434.59, -10.65], p = 0.040). However, prior exposure doses (low versus intermediate) did not significantly differ from each other for pathogen gene expression (low+high vs. intermediate+high: -6.45, 95% CI [-238.65, 225.75], p = 0.997).

**Figure 3.**
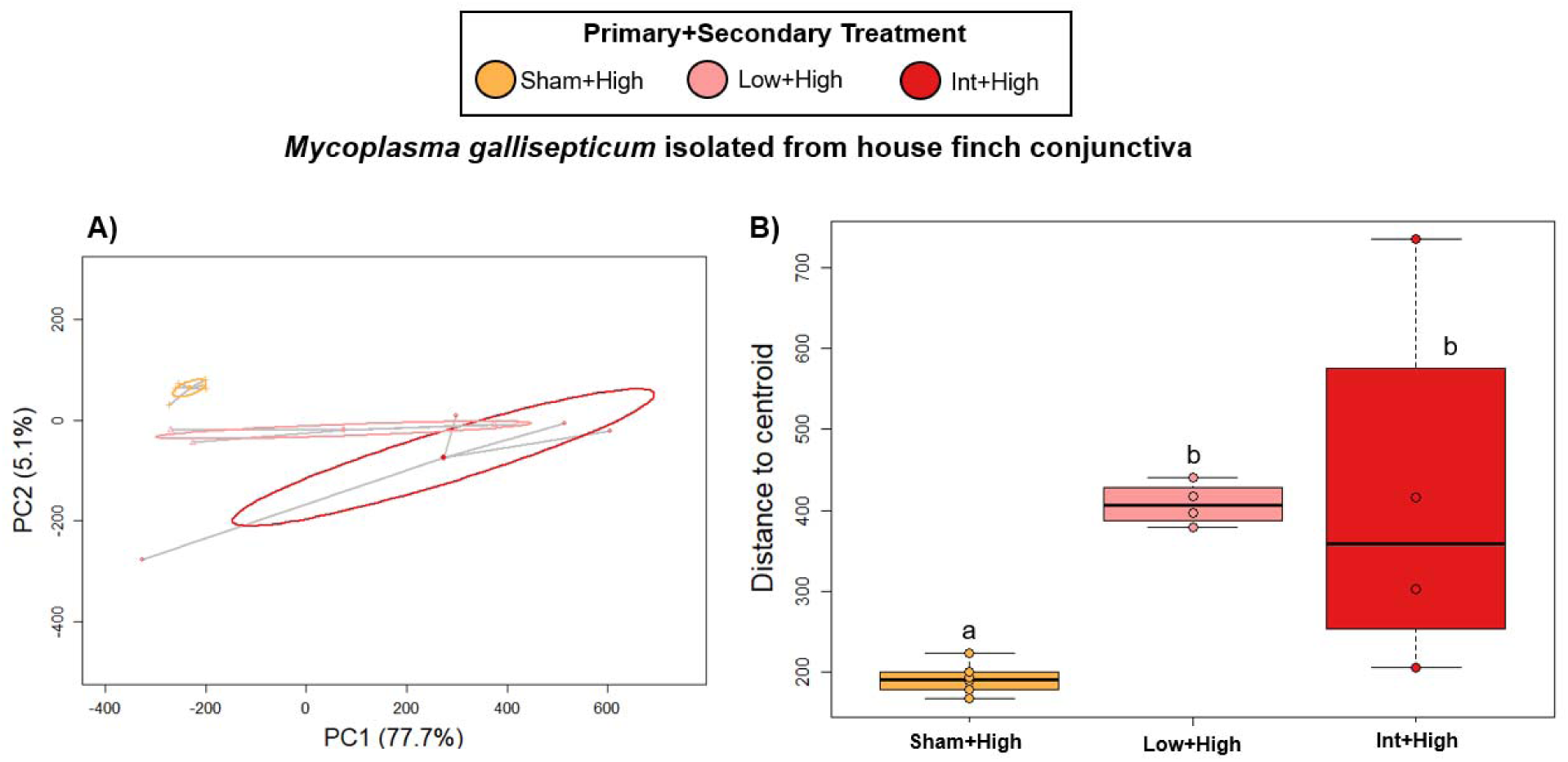
Heterogeneity in gene expression of pathogen (*Mycoplasma gallisepticum*) infecting the conjunctiva of house finches with distinct prior exposure treatments (primary+secondary) three days post-secondary challenge. (A) PCoA plot (principal coordinate analysis) of PERMANOVA (permutational analysis of variance) results using Manhattan distance matrices of r-log-transformed *Mycoplasma gallisepticum* gene expression data. (B) Boxplot of group dispersion values shown with PERMDISP (test for multivariate dispersion homogeneity) results. Significant post-hoc differences are represented by unshared letters between groups. Treatment groups represent the house finch’s treatment that the pathogen was extracted from.

### Functional enrichment analyses

#### Host

When comparing the expression profiles of birds that received a secondary-MG dose, the majority (87/97) of significantly differentially variable (SDV) genes had greater variances in the low+high treatment group relative to those birds that were pathogen-naïve at the time of secondary challenge (low+high > sham+high: 87 genes at FDR 0.1; 55 genes at FDR 0.05). Top SDV genes [full list of SDV genes – Table S5] included two transcript variants of the solute carrier family 28 (*SLC28A2* adj. p = 0.004 and *SLC28A3* adj. p = 0.013); *ALDH1A1* and *BNIP3* (adj. p <0.001), *TNFSF6B* (adj. p = 0.003), *BID* and *GPC1* (adj. p = 0.004); *BATF3* and *ICAM5* (adj. p = 0.006); *NTN4* (adj. p = 0.013); and *IL18BP* (adj. p = 0.014). Two SDV genes (2 at FDR 0.05) had higher variances in birds that had received an intermediate prior dose relative to naïve birds: *MYL2* (adj. p <0.001) and *C1QA* (adj. p = 0.008). Likewise, two SDV genes had greater variance in sham+high over intermediate+high birds: an uncharacterized ncRNA (adj. p < 0.001) and *MUC-5B* (adj. p = 0.003). Six SDV genes were more variable in the sham+high over the low+high birds (2 genes at FDR 0.05: *NUDT7* [adj. p = 0.048] and *SPRR5* [adj. p = 0.040]).

The list of SDV genes that had greater variance in low+high relative to sham+high birds was enriched for nineteen GO terms (Biological Process; Figure 4). Of the 87 genes in the gene list, the over-representation analysis removed 24 with no GO terms that exceeded the FDR threshold (0.05). The most highly significant terms belonged to the GO parent terms “response to stimulus” (GO:0050896), or “biotic stimulus” (GO:0009607). These included “defense responses to other organisms” (GO:0098542) and “biological processes involved in interspecies interaction” (GO:0044419), such as “responses to viruses” (GO:0009615) and “molecules of bacterial origin” (GO:0002237). Multiple terms were enriched for “cellular response to biotic stimulus” (GO:0071216) or “chemical stimulus” (GO:0070887). No significant GO terms were returned for Cellular Components or Molecular Functions. Over-representation analyses were not performed for any other SDV conjunctival gene lists due to their small number of genes.

**Figure 4.**
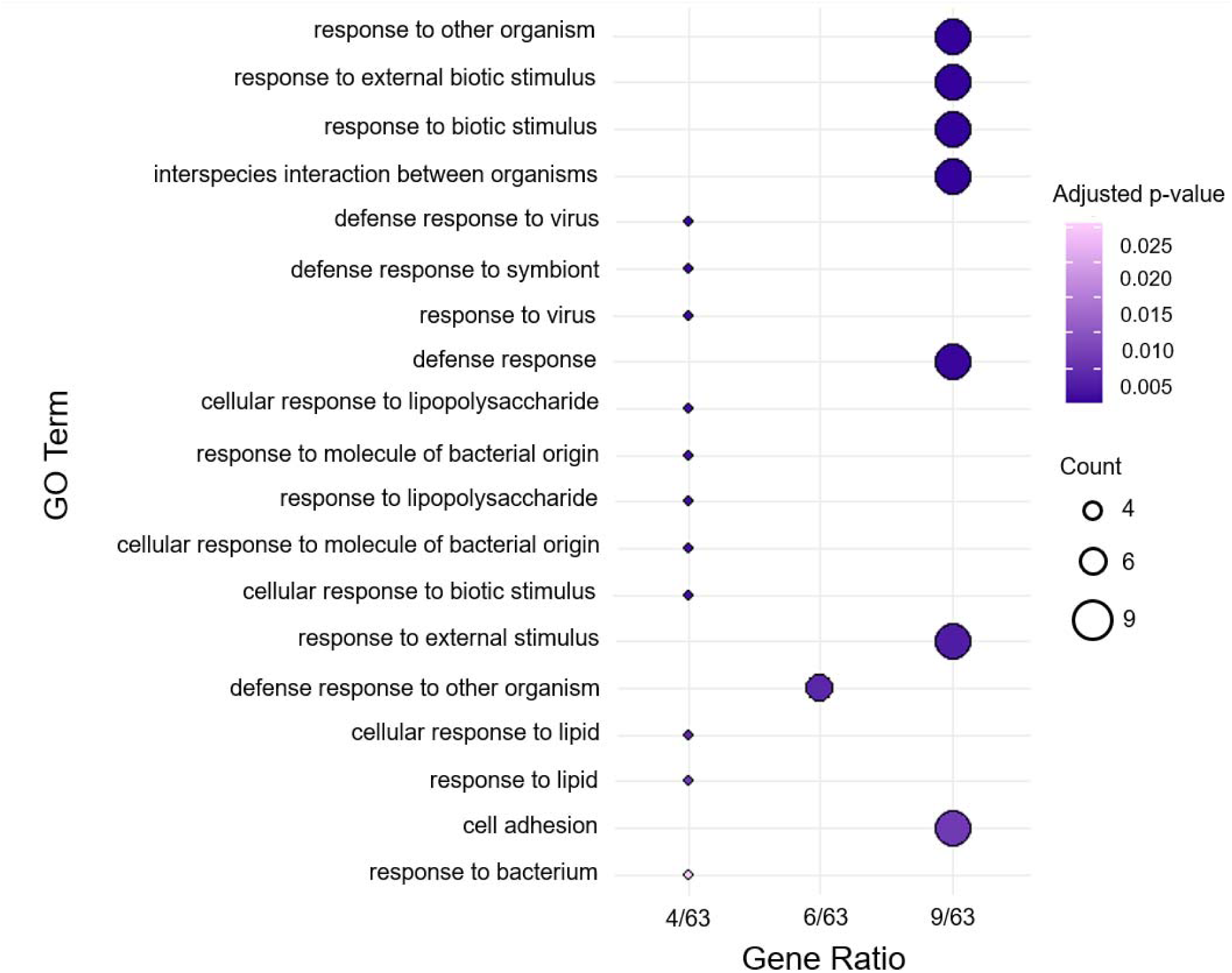
Dotplot of significantly enriched gene ontology (GO) terms for Biological Processes. Terms (n = 19; FDR 0.05) were derived from host genes that were significantly differentially variable (SDV) in the conjunctiva of pathogen-challenged house finches with low dose prior exposure relative to those naïve at the time of challenge. Terms are ordered by ascending Gene Ratio (X-axis; the proportion of genes assigned to that term out of the total considered) and descending adjusted p-value (Y-axis). Dots are sized according to the number of genes in each category and colored by their adjusted p-value (darker colors indicate greater statistical significance).

#### Pathogen

The expression of 104 genes (67 at FDR 0.05) were significantly more variable in MG infecting the conjunctiva of house finches that received a low prior dose (low+high) relative to pathogen infecting birds naïve at the time of challenge (sham+high). Pathogen infecting intermediate+high birds had 3 SVD genes over the sham+high group (3 at FDR 0.05). No genes were more variable in the sham+high over the intermediate+high group, but one (1 at FDR 0.05) was higher in pathogen derived from naïve birds compared to those from low dose prior exposure birds: *upgC* (adj. p = 0.014) [full list of SDV genes - Table S6]. Interestingly, no GO terms were significantly enriched for the 55 genes that were retained in the analysis for the low+high vs sham+high gene list [GO enrichment results - Table S7].

### Pathogen loads

Pathogen loads (log_10_) in eye swabs collected from birds three days post-secondary MG challenge were not significantly different by treatment type in a Kruskal-Wallis test (*X*^2^ = 5.66, df = 2, p = 0.059; Figure 5). Planned comparison Wilcoxon rank sum tests indicated a significant difference between the sham and intermediate prior exposure treatment groups (p = 0.014), but not between sham and low dose prior exposure (p = 0.241). Coefficients of variation (CV) for pathogen loads were highest in eyeswabs collected from birds with low dose prior exposure (CV = 0.71) and lowest for those that had received a sham primary dose (CV = 0.03; intermediate prior exposure CV = 0.47).

**Figure 5.**
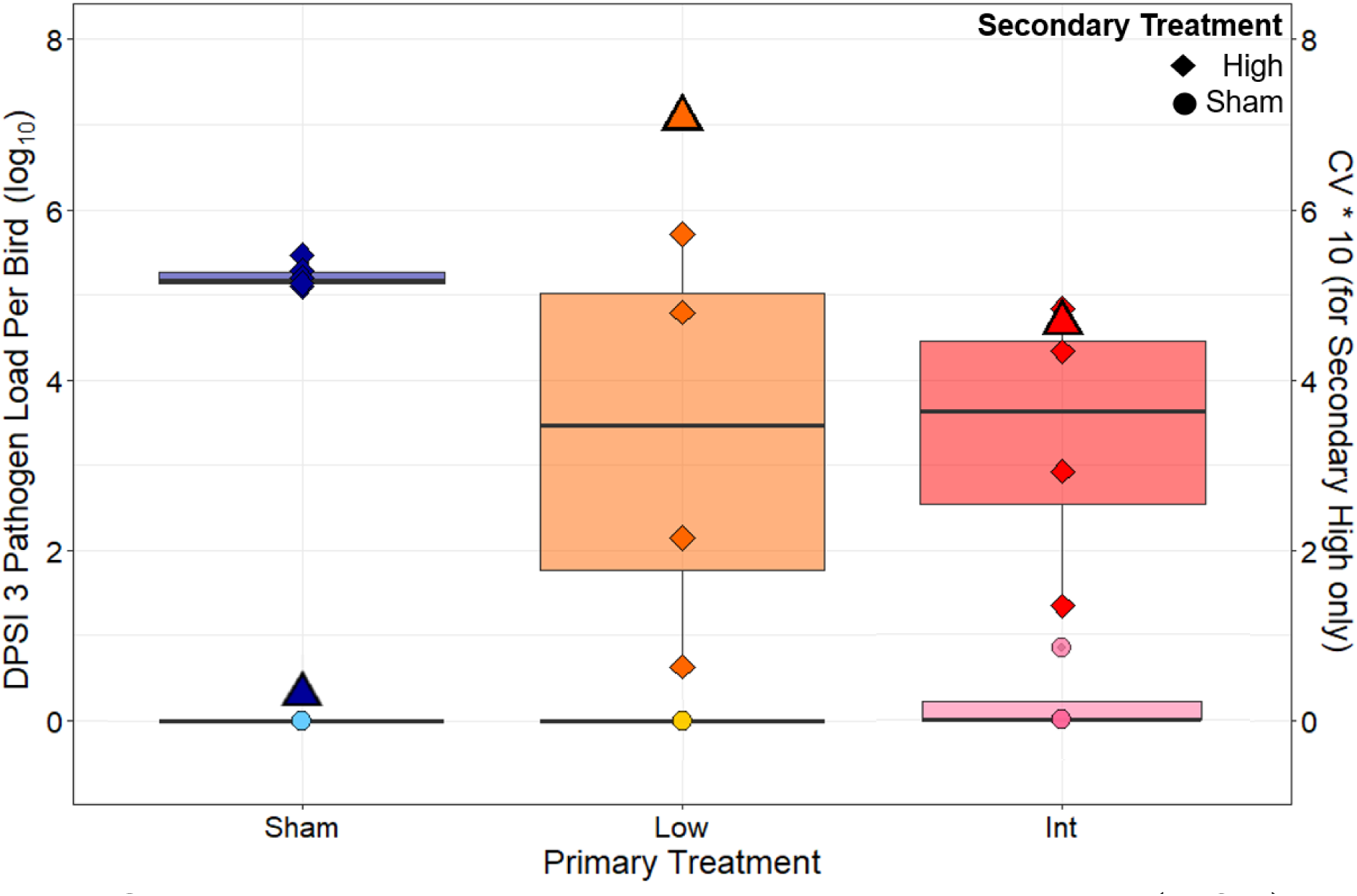
Conjunctival pathogen loads three days post-secondary inoculation (DPSI 3) for individuals from each prior exposure treatment. Loads are on a log_10_ scale. Boxes indicate the interquartile ranges (IQR), encompassing the 25^th^ to 75^th^ percentiles, with the horizontal lines representing the medians (50^th^ percentiles). Boxes and data points are colored to represent the primary treatment (blue: sham; orange: low; red: intermediate) with lighter (sham) or darker (high) shades of each color indicating the secondary treatment. Additionally, data points are shaped according to secondary treatment (circle: sham; diamond: high). The coefficient of variation (CV; scaled by a factor of 10) is shown (triangles) for the high secondary treatment groups only. Data points for birds that received a secondary sham dose are shown but were not included in analyses.

## Discussion

For an infecting pathogen, the within-host environment functions as the “habitat” within which it must balance replication with evasion of the immune system, and from which it must ultimately transmit to a new, susceptible host. Thus, the degree of population-level variation in that within-host environment has significant downstream consequences for the host-pathogen coevolutionary dynamic (Ganz & Ebert, 2010; Gibson & Lively, 2019; Morley et al., 2017; P. S. White et al., 2020). Here we explicitly consider how distinct within-host environments (differing in acquired protection from prior exposure) can alter the plastic nature of gene regulatory responses during infection (Råberg et al., 1998). Specifically, we used a two-part experiment with songbirds exposed to varying doses of an endemic bacterial pathogen to ask whether prior pathogen exposure induces heterogeneity in both host and pathogen gene expression during *in vivo* infection. We found that prior exposure significantly augmented heterogeneity in gene expression in the host tissue that serves as the primary site of infection (palpebral conjunctiva), though not the remote secondary lymphoid tissue tested (spleen). Importantly, pathogen isolated from hosts with prior pathogen exposure exhibited greater gene expression heterogeneity in a manner similar to the host conjunctiva results. Overall, our results are consistent with the hypothesis that prior host exposure induces a within-host environment that promotes heterogeneous gene expression across both hosts and pathogens, with key potential consequences for pathogen evolution.

In a disease system where prior pathogen exposure provides strong and durable acquired protection, exposing a population to a controlled pathogen dose might homogenize its response upon reinfection relative to an unprimed population. However, the immune protection generated by prior pathogen exposures and vaccinations are often incomplete (Le et al., 2021), resulting in heterogeneity in host susceptibility (Gomes et al., 2014; Hawley et al., 2024; Langwig et al., 2017) and pathogen loads (Garrett-Larsen et al., 2025 [Preprint]) upon reinfection/challenge. This, in turn, has implications for pathogen infectiousness (Bailey et al., 2020; Leon et al., 2025; Puhach et al., 2022), a key proxy for pathogen fitness. Our results indicate that this disease-response heterogeneity in hosts with some standing protection extends to gene regulation in the within-host environment: despite receiving an identical challenge, hosts with prior exposure had augmented gene expression heterogeneity during reinfection (in the conjunctiva) over hosts that were pathogen-naive. Further, this heterogeneity was moderately dose-dependent: animals with low-dose prior exposure had greater within-group heterogeneity. Importantly, birds that received a secondary sham control dose (and thus were not actively infected at the time of sampling) did not differ in gene expression heterogeneity regardless of their prior exposure history. This indicates that the detected differences in host heterogeneity occurred in response to a secondary MG challenge and were not ongoing immune responses from initial MG exposure (i.e., these responses were only induced when birds were re-exposed to MG and were not just carryovers from their initial MG exposure). Finally, immune genes were the primary driver of the observed heterogeneity, as revealed by functional enrichment analyses. This supports the prediction that the observed heterogeneity of gene expression induced by prior exposure occurs for genes relevant to the host’s immune response. This is largely unsurprising given that animals were experimentally infected and immune genes are among the most transcriptionally variable in the genome (Wolf et al., 2023). However, these analyses are useful in identifying those genes and GO terms/pathways which have functional significance for prior exposure-induced heterogeneity.

In the conjunctival tissue of the low+high treatment group (the most heterogeneous group), nineteen GO terms (Biological Processes) were SDV-gene enriched relative to sham+high birds. These terms covered various aspects of the immune system’s response to pathogens (e.g., “defense responses to other organisms”). Examination of the function of individual SDV genes lends additional insight into the mechanisms of this response. In the conjunctiva, prior exposure with a low MG dose appeared to diversify expression of the inflammatory immune response of the reinfected birds, which is driven by the heterogeneous responses of macrophages and dendritic cells that interact with proinflammatory T cells. Namely, in the birds pre-stimulated with a low pathogen dose, secondary challenge with MG increased variation in the expression of *BNIP3*, *BATF3*, *TNFSF6B*, and *IL18BP*. *BNIP3* (BCL2) is a pro-apoptotic protein that up-regulates the IL17 (Th17) pathway by increasing phosphorylation of NF-kappa B (Song et al., 2025). Complementary to this action, BATF3-dependent dendritic cells help up-regulate Th1 immune responses through the production of type 1 cytokines such as interferon gamma (Xu et al., 2024). Anti-inflammatory action is provided by IL18BP, which serves as an inhibitor of IL18, an important proinflammatory cytokine (Menachem et al., 2024). Finally, this response could be regulated by the expression of *TNFSF6B*, which belongs to the TNF superfamily, members of which are produced by various leukocytes during inflammation. Similarly, *C1QA*, one of two SDV genes with increased variance in birds that received an intermediate prior exposure dose, is involved in the complement cascade which promotes the killing of pathogens (Mulder et al., 2021). Although birds naïve at the time of secondary challenge (sham+high) were the most homogenous group, two genes had greater variance when compared to the intermediate+high treatment group. One of these was *MUC-5B*, whose protein is expressed into the mucus of the respiratory tract where it supports lung macrophage function (Roy et al., 2014). We can speculate that the relatively higher initial stimulation with the pathogen provides consistently higher protection in the birds upon their secondary contact with MG through increased mucosal protection.

Prior exposure alone broadly induced conjunctival gene expression heterogeneity in birds when re-exposed (Figure 2A), but the degree (i.e., dose) of prior exposure appeared to at least partially underlie the extent of that heterogeneity. Birds that received an intermediate prior exposure dose had increased heterogeneity that statistically overlapped with all other treatment groups, whereas animals that received a low prior exposure dose had significantly elevated heterogeneity relative to birds that were naive to MG at the time of infection (and all sham-secondary treatment groups). Although we can only speculate as to the exact mechanisms underlying the observed increases in heterogeneity, it is reasonable that “vaccination” efficacy or a higher prior exposure dose (here, our intermediate dose) would be more likely to initiate more complete protection that homogenizes downstream expression responses. Indeed, past work suggests that low dose prior exposure leads to more variable protection than higher exposure doses (i.e., as observed through pathogen loads – Leon et al. 2019; Garrett-Larsen et al., 2025 [Preprint]), a result mirrored in the pathogen loads collected in this study. Here, pathogen loads (as measured by CV and sampled three days post-secondary challenge) were most variable when collected from birds with low prior exposure compared to those that received a high prior dose or were naïve at the time of challenge. However, larger sample sizes are required to definitively determine whether exposure dose drives heterogeneity. It is important to understand whether the extent of heterogeneity is dose-dependent because in this disease system, and many others, low doses are more likely to be experienced in nature (Adelman et al., 2013; Dhondt et al., 2007).

Despite clear patterns for the conjunctiva, there were no detectable differences in either gene expression heterogeneity or group central tendencies (i.e., centroids) in the remote secondary lymphoid tissue tested - the spleen. It is possible that the central role of the spleen in the immune system obscured detectable differences: the spleen likely addresses multiple challenges at any given time, including any background symbionts. This could explain why the control birds (sham+sham) were similar to all other treatment groups in terms of splenic gene expression profile (overlapping centroids) and heterogeneity (equivalent dispersion values). Most likely, however, the lack of a splenic response is a result of sample collection timing, just three days post-challenge. Bonneaud et al. (2012) found that splenic gene expression was substantially muted on day 3 relative to later sampling, although this was in the context of an initial MG-infection. Given that animals in our study were reinfected, we would expect some anamnestic response by day 3 and it is therefore surprising that there were no treatment differences. Regardless, at such an early timepoint, it is likely the directly infected tissue, rather than a remote secondary one, that is the primary source of selective pressures on the pathogen.

Pathogen gene expression heterogeneity was also augmented by host prior exposure and largely mirrored the degree of heterogeneity exhibited by the host tissue (conjunctiva) it was infecting. MG extracted from hosts that were naïve at the time of challenge were highly homogenous in their regulatory response, whereas those infecting hosts with any degree of prior exposure had elevated heterogeneity. However, unlike in the host tissue, there was no indication of an effect of prior dose received, and functional enrichment analyses did not identify any overrepresented GO terms. Notable, too, was the absence of *vlhA* expression heterogeneity. Phenotypic variation of phase variable genes like *vlhA* is a common bacterial strategy for rapidly and plastically responding to novel environments (i.e., hosts/tissues) by generating a heterogeneous, rather than a clonal population of bacteria (Parkhill 2008). This has potentially played an important role in MG’s ability to colonize new hosts (Pflaum et al., 2020). For MG, rapid, phase-variable changes in *vlhA* gene expression are associated with the infection of different tissues and host species (Pflaum et al., 2020), and it is possible that this strategy at *vlhA* or other loci contributes to the successful infection of hosts with prior exposure and some degree of immunity (Leon et al., 2019). But, interestingly, no significant heterogeneous *vlhA* expression was seen between birds in our study. This suggests that our pre-exposure doses did not create conditions sufficient to alter *vlhA* expression heterogeneity at this timepoint, or that changes in *vlhA* expression were not detectable via *in vivo* pathogen sequencing.

Overall, we found that prior exposure to MG induced similar patterns of augmented expression heterogeneity for both host and pathogen. However, the directionality and causality of this effect remains unresolved. Differences in pathogen loads – which are confounded with treatment – or MG’s response to hosts with standing protection, could have contributed to the variation in both host and pathogen gene expression. Bacterial gene expression is a response to its immediate microenvironment which includes not just the host’s tissue but also the density, identity, spatial arrangement, and phenotype of nearby bacterial cells (van Vliet et al., 2018). This density-dependent nature in bacterial gene expression could mean that host prior exposure alone augments pathogen heterogeneity by having conferred some degree of host protection that modulates MG’s replication and density. It is also possible that the augmented variability in gene expression within host tissues directly stimulates higher variability in pathogen gene expression, or vice versa. A final explanation for the drivers of gene expression heterogeneity is that the selective pressures on a host vary according to whether it is the primary infection or a reinfection. In this system, the first infection is more severe than subsequent infections – i.e., higher pathogen loads and disease severity (Leon & Hawley, 2017) – which is correlated with reduced anti-predator responses (Adelman et al., 2017) and mortality in the wild (Faustino et al., 2004). If there is an optimal response to primary infection and a greater cost to deviating from it, perhaps we would expect to see population uniformity in that response, such as we do in the naïve birds when challenged for the first time (i.e., sham+high treatment). These possibilities are not mutually exclusive, and our study design cannot determine their relative contributions. Regardless, prior exposure history induced parallel patterns of regulatory heterogeneity in both host and pathogen, which suggests that, beyond genetic variation, gene expression is key to our understanding of reciprocal selection pressures between hosts and pathogens.

In summary, we found that prior pathogen exposure generated gene expression heterogeneity in both a songbird host and its infecting bacterial pathogen. That is, the same ecological condition drove phenotypic variation in both organisms. We also found that lower dose prior exposure produced the greatest degree of heterogeneity in the host. This broadly correlates with the dose-dependent heterogeneity observed in other traits, namely pathogen load, in this and other studies (Leon et al. 2019; Hawley et al. 2024; Garrett-Larsen et al., 2025 [Preprint]). If, mechanistically, variation in gene expression influences disease trait phenotypes such as host susceptibility, tolerance (i.e., per unit pathogen effect on host fitness – Råberg et al. 2009), or competence (i.e., ability to infect another susceptible host – Gervasi et al., 2015), then this diversity generated by ecological condition can directly influence epidemiological outcomes and, ultimately, host/pathogen fitness and evolution, including strain differentiation and virulence evolution. Overall, our results demonstrate that, beyond underlying genetic diversity, gene expression should be accounted for when considering the reciprocal costs and benefits of host-pathogen diversity and overall trajectory of the coevolutionary relationship.

## Acknowledgements

This research was made possible by grant R01GM144972 from the National Institutes of Health (NIH) Ecology and Evolution of Infectious Diseases (EEID) program as well as the Inter-ACTION project No. LUAUS24184 supported by the Ministry of Education, Youth and Sports of the Czech Republic. All birds were handled and captured under U.S. Fish and Wildlife Service (MB154804-0) and Virginia Department of Game and Inland Fisheries (066646) permits. All housing and experimental protocols were approved by the Virginia Tech Institutional Animal Care and Use Committee (IACUC). All the research complies with applicable laws on sampling from natural populations and animal experimentation (including the ARRIVE guidelines). We thank our collaborators on the NIH-EEID grant for their valuable discussions about disease trait heterogeneity: Lauren Childs, Arietta Fleming-Davies, and Katie Talbott. Many thanks to the Virginia Tech undergraduate and graduate students who helped to catch and quarantine the birds, and collect data and samples (Madeline Alt, Alicia Arneson, Annabel Coyle, Zhang Gao, Noelle Hodges, Marissa Langager, Riley Meyers, Sara Teemer, and Caro Vela).

