## Supplementary for "Host prior exposure augments heterogeneity in gene expression in both host and pathogen during *in vivo* infection"

**Supplemental Methods**

**Table S1.** Experimental timeline. Sample types and their collection timepoints, with associated dates, across the experiment for all birds. Timepoints are reported relative to days of inoculation: DPPI (days post-primary inoculation) and DPSI (days post-secondary inoculation), with “X” specifying the sample types collected on each occasion. The horizontal dotted line indicates the transition from primary to secondary portions of the study.

| **Time Points** | | **Samples Collected** | | | | |
| --- | --- | --- | --- | --- | --- | --- |
| **Date** | **Sample Timepoint** | **Swabs** | **Scores** | **Plasma** | **Tissues** | **Inoculate** |
| 8/29/2022 | **DPPI -4** | X |  | X |  |  |
| 9/2/2022 | **DPPI 0** |  |  |  |  | X |
| 9/9/2022 | **DPPI 7** | X | X |  |  |  |
| 9/16/2022 | **DPPI 14** | X | X |  |  |  |
| 9/23/2022 | **DPPI 21** |  | X |  |  |  |
| 9/30/2022 | **DPPI 28** |  | X |  |  |  |
| 10/7/2022 | **DPPI 35** |  | X | X |  |  |
| 10/14/2022 | **DPPI 42 / DPSI 0** | X | X |  |  | X |
| 10/17/2022 | **DPSI 3** | X | X |  | X |  |

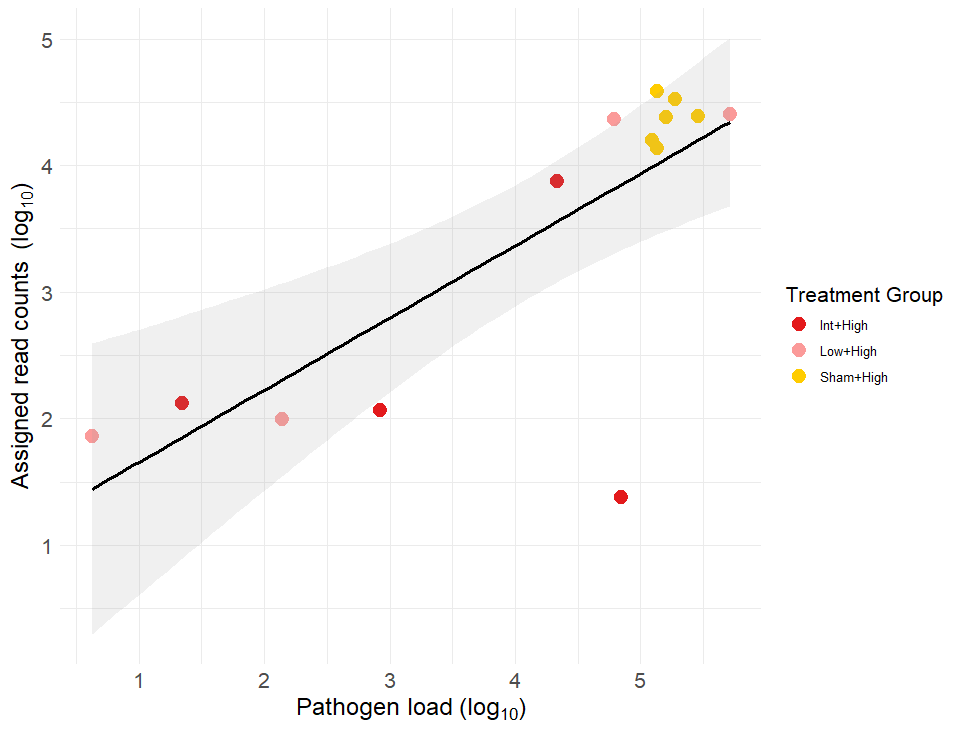

**Figure S1.** Number of reads assigned to *Mycoplasma gallisepticum* genes (i.e. counts) relative to that sample’s pathogen load (logged scales). Data are shown only for samples that received a secondary pathogen challenge. Sample datapoints are colored by treatment group.

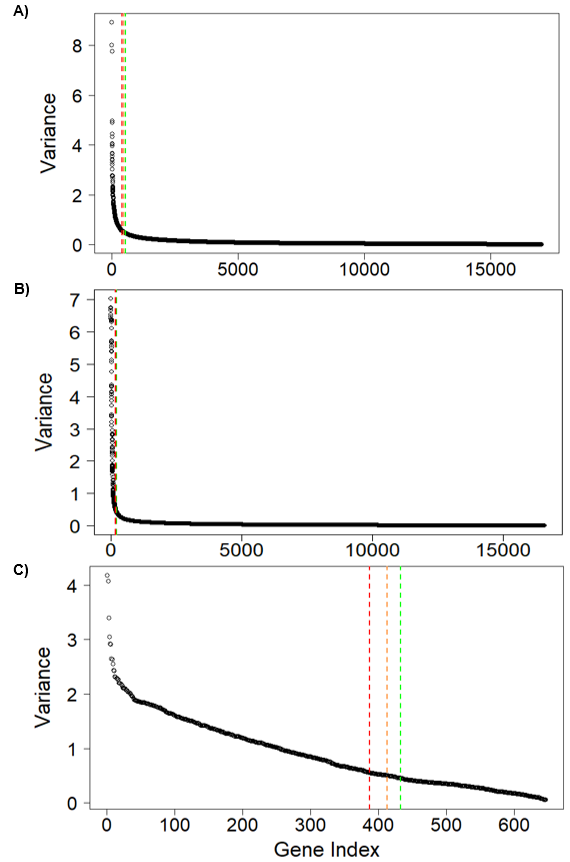

**Figure S2.** Genes (gene index) ordered by descending expression variance for three different sample types (A: house finch conjunctiva; B: house finch spleen; C: infecting *Mycoplasma gallisepticum*). Vertical lines indicate the gene index where the variance is greater than 0.55 (green), 0.50 (orange), and 0.45 (red).

*DNA extractions and qPCR assays*

Eyeswab samples were kept on ice and stored at -20°C until nucleic acid extraction with 96-well DNeasy Blood and Tissue Kits (Qiagen, Hilden, Germany). Extracted swab samples were assayed via qPCR (QuantiNova Kit [Qiagen, Hilden, Germany]) with probes and primers designed to target the *mgc2* gene (Grodio et al., 2008; Hawley et al., 2011). Assays were performed on a QuantStudio 5 machine using manufacturer-specified cycling conditions. Estimates of MG pathogen copies per sample were derived from standard curves of 1:10 serially diluted (3.86 x 10^1^ – 3.86 x 10^8^ copies) pCR4-TOPO plasmids containing an *mgc2* insert. Three negative controls were included per extraction and qPCR plate, and standards were run in triplicate for each qPCR assay.

**Supplemental Results**

*Assigned reads*

An average of 7.30M ± 0.29 SE (range : 2.08 – 9.11) and 6.57M ± 0.15 SE (range: 5.15 – 8.54) reads were assigned to genes (i.e., counts) for conjunctiva and spleen samples, respectively. Sequencing depth for MG samples was contingent on the pathogen load of infected birds, which was dependent on the bird’s prior treatment (Figure S1) and therefore ranged widely. MG samples had 14,760 ± 3,639.81 SE (range: 24 – 38,810) reads assigned to genes, where samples from sham+high birds had typically the greatest number of reads and samples from birds with some degree of prior exposure had lower read assignments.

**Table S2.** PERMANOVA and PERMDISP results (F-value and p-value) for conjunctiva, spleen, and MG tissue using different variance cutoffs (± 10% of 0.5). “Genes retained” refers to how many genes were kept for the analysis after filtering with a given variance threshold. Bolded values are statistically significant.

| **Variance threshold** | **0.45** | | | | | **0.55** | | | | |
| --- | --- | --- | --- | --- | --- | --- | --- | --- | --- | --- |
|  | **PERMANOVA** | | **PERMDISP** | | **Genes retained** | **PERMANOVA** | | **PERMDISP** | | **Genes retained** |
| **Tissue** | **F** | **p** | **F** | **p** |  | **F** | **p** | **F** | **p** |  |
| Conjunctiva | 6.24 | **<0.001** | 5.97 | **0.003** | 544 | 5.98 | **<0.001** | 6.35 | **<0.001** | 394 |
| Spleen | 0.488 | 0.932 | 0.802 | 0.588 | 199 | 0.460 | 0.948 | 0.784 | 0.557 | 162 |
| MG | 3.06 | **0.05** | 5.58 | **<0.001** | 433 | 3.06 | **0.029** | 5.48 | **<0.001** | 387 |

**Table S3.** P-values from Tukey HSD post-hoc comparisons of PERMDISP results when using different variance cutoffs (± 10% of 0.5) for conjunctiva, spleen, and MG tissues. Bolded values are statistically significant.

| **Variance threshold** | **0.45** | | | **0.55** | | |
| --- | --- | --- | --- | --- | --- | --- |
| **Treatment comparisons** | **Conjunctiva** | **Spleen** | **MG** | **Conjunctiva** | **Spleen** | **MG** |
| high_sham-high_high | 0.788 | 0.820 |  | 0.874 | 0.766 |  |
| low_high-high_high | 0.375 | 0.598 | 0.994 | 0.218 | 0.608 | 0.998 |
| low_sham-high_high | 0.307 | 0.980 |  | 0.327 | 0.969 |  |
| sham_high-high_high | 0.166 | 0.587 | **0.039** | 0.222 | 0.594 | **0.042** |
| sham_sham-high_high | 0.352 | 0.671 |  | 0.514 | 0.666 |  |
| low_high-high_sham | 0.036 | 0.999 |  | 0.025 | 1.000 |  |
| low_sham-high_sham | 0.954 | 0.995 |  | 0.916 | 0.993 |  |
| sham_high-high_sham | 0.876 | 1.000 |  | 0.863 | 1.000 |  |
| sham_sham-high_sham | 0.981 | 1.000 |  | 0.991 | 1.000 |  |
| low_sham-low_high | **0.005** | 0.938 |  | **0.003** | 0.959 |  |
| sham_high-low_high | **0.001** | 1.000 | **0.047** | **0.001** | 1.000 | **0.047** |
| sham_sham-low_high | **0.005** | 1.000 |  | **0.004** | 1.000 |  |
| sham_high-low_sham | 1.000 | 0.950 |  | 1.000 | 0.968 |  |
| sham_sham-low_sham | 1.000 | 0.972 |  | 0.997 | 0.981 |  |
| sham_sham-sham_high | 0.998 | 1.000 |  | 0.993 | 1.000 |  |

**Table S4.** Alignment results from NCBI’s Nucleotide BLAST database for unannotated loci (locus tag; organized by ascending alphanumeric order) that were significantly differentially variable in Brown-Forsythe tests (statistics in Table S5). Loci are from house finch (*H. mexicanus*) conjunctival tissue. For each locus, the top alignment is reported with its predicted mRNA identity and function (Sequence Description) and the species the sequence belongs to. Top alignments were determined by the percentage of the query sequence that aligned to a database sequence (Query Coverage), the percentage of matching nucleotides within that alignment (Percent Identity), and the number of random sequences in the database that would be expected to align equally as well (E-value). More than one top alignment is reported per locus if mapping was uncharacterized or ambiguous (i.e., multiple top results of equivalent quality). Genomic Coordinates of the query sequence (in base pairs), nucleotide length of the alignment (Accession Length), and NCBI’s unique identifier (Accession) are reported per locus/aligned sequence.

| **Locus tag** | **Genomic coordinates** | **Sequence Description** | **Scientific Name** | **Query Cover** | **E value** | **Per. ident** | **Acc. Len** | **Accession** |
| --- | --- | --- | --- | --- | --- | --- | --- | --- |
| LOC132324463 | 4943355-4953837 | uncharacterized transcript variant, misc_RNA | *H. mexicanus* | 38% | 0 | 100 | 3976 | [XR_009485629.1](https://www.ncbi.nlm.nih.gov/nucleotide/XR_009485629.1?report=genbank&log$=nucltop&blast_rank=1&RID=UCM8X70D016) |
|  |  | anaphase promoting complex subunit 1 (ANAPC1), mRNA | *H. mexicanus* | 10% | 2.00E-142 | 100 | 6142 | [XM_059840580.1](https://www.ncbi.nlm.nih.gov/nucleotide/XM_059840580.1?report=genbank&log$=nucltop&blast_rank=18&RID=UCM8X70D016) |
| LOC132324745 | 16472965-16489410 | uncharacterized transcript variant, mRNA | *H. mexicanus* | 69% | 0 | 100 | 11409 | [XM_059841211.1](https://www.ncbi.nlm.nih.gov/nucleotide/XM_059841211.1?report=genbank&log$=nucltop&blast_rank=1&RID=UCMDNGHS016) |
|  |  | mycocerosic acid synthase-like, mRNA | *Serinus canaria* | 41% | 0 | 97.19 | 6755 | [XM_018909287.3](https://www.ncbi.nlm.nih.gov/nucleotide/XM_018909287.3?report=genbank&log$=nucltop&blast_rank=4&RID=UCMDNGHS016) |
| LOC132325633 | 116913226-116936891 | interferon-induced very large GTPase 1-like, mRNA | *H. mexicanus* | 100% | 0 | 100 | 9619 | [XM_059843171.1](https://www.ncbi.nlm.nih.gov/nucleotide/XM_059843171.1?report=genbank&log$=nucltop&blast_rank=1&RID=7NB1B0PG015) |
| LOC132325636 | 117212349-117237099 |  |  | 100% | 0 | 100 | 88864 | [XM_059843173.1](https://www.ncbi.nlm.nih.gov/nucleotide/XM_059843173.1?report=genbank&log$=nucltop&blast_rank=1&RID=7NBEXMRC015) |
| LOC132325944 | 878649-881975 | interleukin-8-like, mRNA | *H. mexicanus* | 37% | 0 | 100 | 1229 | [XM_059843479.1](https://www.ncbi.nlm.nih.gov/nucleotide/XM_059843479.1?report=genbank&log$=nucltop&blast_rank=2&RID=UCMJKM27016) |
| LOC132326811 | 75649047-75651185 | BCL2/adenovirus E1B 19 kDa protein-interacting protein 3-like, transcript variant, mRNA | *Serinus canaria* | 66% | 0 | 88.86 | 1417 | [XM_018923151.3](https://www.ncbi.nlm.nih.gov/nucleotide/XM_018923151.3?report=genbank&log$=nucltop&blast_rank=2&RID=UCMS8Z2E016) |
| LOC132328383 | 66776387-66777610 | uncharacterized, transcript variant, ncRNA | *H. mexicanus* | 100% | 0 | 100 | 1052 | [XR_009486767.1](https://www.ncbi.nlm.nih.gov/nucleotide/XR_009486767.1?report=genbank&log$=nucltop&blast_rank=1&RID=7NBR3367015) |
| LOC132328387 | 19269610-19306616 | mucin-5B-like, mRNA | *H. mexicanus* | 37% | 0 | 100 | 13568 | [XM_059848206.1](https://www.ncbi.nlm.nih.gov/nucleotide/XM_059848206.1?report=genbank&log$=nucltop&blast_rank=1&RID=UCMZVH70013) |
| LOC132329864 | 22998759-23004477 | cytochrome P450 2C9-like, mRNA | *H. mexicanus* | 100% | 0 | 100 | 1742 | [XM_059851441.1](https://www.ncbi.nlm.nih.gov/nucleotide/XM_059851441.1?report=genbank&log$=nucltop&blast_rank=1&RID=7NBWPUGH014) |
| LOC132329926 | 25834586-25839582 | interferon-induced protein with tetratricopeptide repeats 5-like, mRNA | *H. mexicanus* | 43% | 0 | 100 | 2169 | [XM_059851583.1](https://www.ncbi.nlm.nih.gov/nucleotide/XM_059851583.1?report=genbank&log$=nucltop&blast_rank=1&RID=UCNYCHJF016) |
| LOC132333113 | 5739967-5748563 | sodium/nucleoside cotransporter 2-like | *H. mexicanus* | 100% | 100 | 100 | 2231 | [XM_059857890.1](https://www.ncbi.nlm.nih.gov/nucleotide/XM_059857890.1?report=genbank&log$=nucltop&blast_rank=1&RID=7NCKC2DK014) |
| LOC132331999 | 14912353-14969826 | IgGFc-binding protein-like, mRNA | *H. mexicanus* | 31% | 0 | 100 | 17519 | [XM_059855990.1](https://www.ncbi.nlm.nih.gov/nucleotide/XM_059855990.1?report=genbank&log$=nucltop&blast_rank=8&RID=UCP1TJRB016) |
| LOC132332265 | 10104726-10106972 | 2'-5'-oligoadenylate synthase 1-like, mRNA | *H. mexicanus* | 78% | 0 | 100 | 1737 | [XM_059856358.1](https://www.ncbi.nlm.nih.gov/nucleotide/XM_059856358.1?report=genbank&log$=nucltop&blast_rank=2&RID=UCP5RDS5016) |
| LOC132332352 | 14296566-14341553 | netrin-4-like, mRNA | *H. mexicanus* | 100% | 0 | 100 | 4603 | [XM_059856557.1](https://www.ncbi.nlm.nih.gov/nucleotide/XM_059856557.1?report=genbank&log$=nucltop&blast_rank=1&RID=7NC1DCY0014) |
| LOC132332733 | 15816802-15823504 | C-signal-like, transcript variant, mRNA | *H. mexicanus* | 27% | 1.00E-102 | 100 | 935 | [XM_059857306.1](https://www.ncbi.nlm.nih.gov/nucleotide/XM_059857306.1?report=genbank&log$=nucltop&blast_rank=47&RID=UCP9M1B9016) |
| LOC132332991 | 3099884-3105736 | peroxisomal coenzyme A diphosphatase NUDT7-like, mRNA | *H. mexicanus* | 26% | 0 | 100 | 1538 | [XM_059857721.1](https://www.ncbi.nlm.nih.gov/nucleotide/XM_059857721.1?report=genbank&log$=nucltop&blast_rank=2&RID=UCRJK69P013) |
| LOC132332995 | 3135607-3141459 |  |  |  |  |  |  |  |
| LOC132333048 | 3498788-3508238 | uncharacterized, transcript variant, ncRNA | *H. mexicanus* | 46% | 0 | 98.98 | 12733 | [XR_009488043.1](https://www.ncbi.nlm.nih.gov/nucleotide/XR_009488043.1?report=genbank&log$=nucltop&blast_rank=3&RID=UCRT94PW016) |
|  |  | fatty acyl-CoA hydrolase precursor, medium chain-like (LOC132333048), transcript variant X2, mRNA | *H. mexicanus* | 25% | 0 | 100 | 2319 | [XM_059857782.1](https://www.ncbi.nlm.nih.gov/nucleotide/XM_059857782.1?report=genbank&log$=nucltop&blast_rank=16&RID=UCRT94PW016) |
| LOC132334445 | 4794083-4816395 | uncharacterized, mRNA | *H. mexicanus* | 13% | 0 | 100 | 2906 | [XM_059860518.1](https://www.ncbi.nlm.nih.gov/nucleotide/XM_059860518.1?report=genbank&log$=nucltop&blast_rank=14&RID=UCRW9HJ7016) |
|  |  | hepatitis A virus cellular receptor 1 homolog, transcript variant, mRNA | *Passer domesticus* | 9% | 0 | 90.74 | 3238 | [XM_064387754.1](https://www.ncbi.nlm.nih.gov/nucleotide/XM_064387754.1?report=genbank&log$=nucltop&blast_rank=1&RID=UCRW9HJ7016) |
| LOC132335646 | 6854890-6855705 | waprin-Enh1-like, mRNA | *H. mexicanus* | 62% | 4.00E-79 | 100 | 488 | [XM_059862431.1](https://www.ncbi.nlm.nih.gov/nucleotide/XM_059862431.1?report=genbank&log$=nucltop&blast_rank=1&RID=UCRZ5VZX013) |
| LOC132335743 | 8272726-8280242 | BPI fold-containing family B member 3-like, transcript variant, mRNA | *H. mexicanus* | 32% | 0 | 100 | 2313 | [XM_059862584.1](https://www.ncbi.nlm.nih.gov/nucleotide/XM_059862584.1?report=genbank&log$=nucltop&blast_rank=1&RID=UCS1VYDJ016) |
| LOC132336561 | 11859483-11862037 | uncharacterized, transcript variant, mRNA | *H. mexicanus* | 58% | 0 | 100 | 1448 | [XM_059864203.1](https://www.ncbi.nlm.nih.gov/nucleotide/XM_059864203.1?report=genbank&log$=nucltop&blast_rank=5&RID=UCS54JRK016) |
| LOC132337150 | 10625257-10634377 | ectonucleoside triphosphate diphosphohydrolase 8-like, transcript variant, mRNA | *H. mexicanus* | 27% | 0 | 100 | 2411 | [XM_059865415.1](https://www.ncbi.nlm.nih.gov/nucleotide/XM_059865415.1?report=genbank&log$=nucltop&blast_rank=1&RID=UCS82H6P013) |
| LOC132337221 | 6080015-6082370 | alpha-1-acid glycoprotein 1-like, transcript variant, mRNA | *H. mexicanus* | 35% | 6.00E-117 | 100 | 814 | [XM_059865571.1](https://www.ncbi.nlm.nih.gov/nucleotide/XM_059865571.1?report=genbank&log$=nucltop&blast_rank=1&RID=UCSBNJWP013) |
| LOC132338534 | 1758685-1759422 | uncharacterized, transcript variant, ncRNA | *H. mexicanus* | 51% | 3.00E-164 | 100 | 554 | [XR_009489466.1](https://www.ncbi.nlm.nih.gov/nucleotide/XR_009489466.1?report=genbank&log$=nucltop&blast_rank=1&RID=UCSEFV7G016) |
| LOC132340016 | 4262201-4262287 | actin gamma-enteric smooth muscle, mRNA | *H. mexicanus* | 100% | 0 | 100 | 1404 | XM_059870726.1 |
| LOC132340235 | 3730629-3733539 | protein MRP-126-like, mRNA | *H. mexicanus* | 21% | 8.00E-156 | 100 | 618 | [XM_059871103.1](https://www.ncbi.nlm.nih.gov/nucleotide/XM_059871103.1?report=genbank&log$=nucltop&blast_rank=1&RID=UCSRRRYK013) |
| LOC132340237 | 3774043-3775322 | putative small proline-rich protein 5, mRNA | *H. mexicanus* | 100% | 0 | 100 | 792 | XM_059871105.1 |
| LOC132340793 | 2581604-2584614 | interferon-induced protein with tetratricopeptide repeats 1-like, mRNA | *H. mexicanus* | 72% | 0 | 100 | 2179 | [XM_059871893.1](https://www.ncbi.nlm.nih.gov/nucleotide/XM_059871893.1?report=genbank&log$=nucltop&blast_rank=1&RID=UCSVPTH4016) |
| LOC132340963 | 399554-404029 | intercellular adhesion molecule 5 (ICAM5), mRNA | *H. mexicanus* | 100% | 0 | 100 | 3268 | XM_059872254.1 |
| LOC132341970 | 80069271-80070715 | avidin-like, mRNA | *H. mexicanus* | 45% | 4.00E-106 | 100 | 638 | [XM_059874142.1](https://www.ncbi.nlm.nih.gov/nucleotide/XM_059874142.1?report=genbank&log$=nucltop&blast_rank=17&RID=UCSYAUBW013) |
| LOC132341745 | 61288716-61301172 | programmed cell death 1 ligand 1-like, transcript variant, mRNA | *H. mexicanus* | 100 | 0 | 100 | 2372 | XM_059873756.1 |
| LOC132341936 | 2117299-2139720 | aldehyde dehyrodgenase 1A1, mRNA | *H. mexicanus* | 100 | 0 | 100 | 2013 | XM_059874088.1 |

**Table S5.** Significantly differentially variable genes for birds that received a secondary MG dose, where comparisons are between the sham+high treatment group versus the low+high or intermediate+high treatment groups. Loci are organized by ascending p-value/adjusted p-value and are listed if they had an adjusted p-value of <0.1. Data is grouped according to whether the variance was greater in the sham or prior exposed treatment (“Treatment Comparison”). Variance per gene per treatment group are reported unless a group was not involved in the statistical comparison (“NA”). Alignments for unannotated loci (“LOC”) can be found in Table S4.

| **Treatment Comparison** | **Locus** | **P-value** | **Adjusted P-value** | **Sham+High Variance** | **Low+High Variance** | **Int+High Variance** |
| --- | --- | --- | --- | --- | --- | --- |
| **low+high > sham+high** | LOC132341936 | <0.00001 | 0.00010 | 0.00001 | 0.45360 | NA |
|  | LOC132326811 | 0.00000 | 0.00067 | 0.01188 | 1.63516 | NA |
|  | TNFRSF6B | 0.00002 | 0.00349 | 0.04810 | 4.79841 | NA |
|  | BID | 0.00003 | 0.00403 | 0.02629 | 0.85063 | NA |
|  | LOC132333113 | 0.00005 | 0.00437 | 0.04112 | 1.00862 | NA |
|  | GPC1 | 0.00006 | 0.00437 | 0.06680 | 0.99583 | NA |
|  | BATF3 | 0.00009 | 0.00587 | 0.02089 | 0.96952 | NA |
|  | LOC132340963 | 0.00010 | 0.00601 | 0.01425 | 0.22616 | NA |
|  | LOC132334445 | 0.00032 | 0.01345 | 0.06458 | 0.70963 | NA |
|  | LOC132332352 | 0.00035 | 0.01345 | 0.07398 | 1.02919 | NA |
|  | SLC28A3 | 0.00032 | 0.01345 | 0.04710 | 0.98349 | NA |
|  | GREM2 | 0.00028 | 0.01345 | 0.04864 | 0.88124 | NA |
|  | IL18BP | 0.00039 | 0.01374 | 0.01239 | 2.26146 | NA |
|  | LOC132337221 | 0.00044 | 0.01455 | 0.58597 | 9.58345 | NA |
|  | WIF1 | 0.00060 | 0.01842 | 0.10903 | 1.05438 | NA |
|  | B4GALNT4 | 0.00080 | 0.01959 | 0.06020 | 1.38495 | NA |
|  | NCF1 | 0.00071 | 0.01959 | 0.05514 | 0.45059 | NA |
|  | RTEL1 | 0.00078 | 0.01959 | 0.07220 | 0.97531 | NA |
|  | PLXDC1 | 0.00081 | 0.01959 | 0.03081 | 0.42833 | NA |
|  | LOC132340235 | 0.00092 | 0.02129 | 0.09075 | 2.62717 | NA |
|  | NOXO1 | 0.00116 | 0.02183 | 0.83945 | 4.41105 | NA |
|  | LOC132325944 | 0.00112 | 0.02183 | 0.13561 | 2.39903 | NA |
|  | SLC26A6 | 0.00116 | 0.02183 | 0.03008 | 0.61102 | NA |
|  | BAK1 | 0.00109 | 0.02183 | 0.04674 | 0.51882 | NA |
|  | LOC132341745 | 0.00118 | 0.02183 | 0.05995 | 0.41255 | NA |
|  | LOC132331999 | 0.00131 | 0.02287 | 0.00058 | 0.33865 | NA |
|  | SRPX | 0.00134 | 0.02287 | 0.07047 | 0.89311 | NA |
|  | IL1B | 0.00160 | 0.02485 | 0.21295 | 2.15899 | NA |
|  | ASS1 | 0.00158 | 0.02485 | 0.03555 | 1.20822 | NA |
|  | CDK5RAP1 | 0.00161 | 0.02485 | 0.01232 | 1.29496 | NA |
|  | LOC132341970 | 0.00169 | 0.02521 | 0.18687 | 5.89553 | NA |
|  | ROS1 | 0.00198 | 0.02864 | 0.20705 | 2.05933 | NA |
|  | SLC16A3 | 0.00226 | 0.03168 | 0.04655 | 0.37112 | NA |
|  | DUSP15 | 0.00276 | 0.03754 | 0.00779 | 0.41217 | NA |
|  | TNFRSF4 | 0.00325 | 0.03989 | 0.04823 | 0.75837 | NA |
|  | CLEC3B | 0.00328 | 0.03989 | 0.09517 | 0.39677 | NA |
|  | IGFBP7 | 0.00324 | 0.03989 | 0.04982 | 0.87425 | NA |
|  | LOC132340016 | 0.00351 | 0.04155 | 0.05867 | 0.62993 | NA |
|  | ZC3H12A | 0.00365 | 0.04211 | 0.02034 | 0.91007 | NA |
|  | IGDCC3 | 0.00382 | 0.04310 | 0.08562 | 0.64366 | NA |
|  | HIF1A | 0.00402 | 0.04422 | 0.06038 | 0.37590 | NA |
|  | ATP12A | 0.00429 | 0.04577 | 0.23442 | 3.02805 | NA |
|  | LOC132332265 | 0.00456 | 0.04577 | 0.09265 | 1.70139 | NA |
|  | SRGN | 0.00450 | 0.04577 | 0.05557 | 0.56577 | NA |
|  | SMIM33 | 0.00451 | 0.04577 | 0.02622 | 0.57531 | NA |
|  | LOC132324745 | 0.00490 | 0.04757 | 0.07654 | 2.49713 | NA |
|  | LOC132336561 | 0.00515 | 0.04757 | 0.01463 | 1.11776 | NA |
|  | PPDPFL | 0.00512 | 0.04757 | 0.13240 | 1.24320 | NA |
|  | GPNMB | 0.00570 | 0.04931 | 0.06474 | 1.15833 | NA |
|  | MECR | 0.00550 | 0.04931 | 0.01166 | 0.41491 | NA |
|  | CLDN5 | 0.00564 | 0.04931 | 0.04881 | 0.85947 | NA |
|  | MYLK | 0.00576 | 0.04931 | 0.06998 | 0.83160 | NA |
|  | MMP7 | 0.00604 | 0.04942 | 0.13871 | 3.44185 | NA |
|  | SERPIND1 | 0.00610 | 0.04942 | 0.18655 | 2.12093 | NA |
|  | HAPLN3 | 0.00604 | 0.04942 | 0.11233 | 1.05978 | NA |
|  | LOC132340793 | 0.00700 | 0.05055 | 0.16073 | 1.69713 | NA |
|  | C2H21orf62 | 0.00654 | 0.05055 | 0.06777 | 0.81463 | NA |
|  | MGLL | 0.00676 | 0.05055 | 0.12737 | 1.44560 | NA |
|  | GALNT15 | 0.00643 | 0.05055 | 0.07195 | 0.91826 | NA |
|  | IL1R2 | 0.00683 | 0.05055 | 0.08315 | 0.58533 | NA |
|  | LOC132337150 | 0.00736 | 0.05229 | 0.04168 | 1.88201 | NA |
|  | MME | 0.00781 | 0.05466 | 0.08450 | 0.77587 | NA |
|  | OLFM4 | 0.00851 | 0.05866 | 1.30062 | 9.54059 | NA |
|  | TNIP3 | 0.00866 | 0.05886 | 0.36658 | 1.75175 | NA |
|  | HCK | 0.00976 | 0.06534 | 0.03625 | 0.28352 | NA |
|  | JCHAIN | 0.00999 | 0.06595 | 0.04465 | 2.80494 | NA |
|  | LOC132329864 | 0.01054 | 0.06856 | 0.00715 | 0.07475 | NA |
|  | STING1 | 0.01111 | 0.07130 | 0.03225 | 0.45772 | NA |
|  | OTOP3 | 0.01152 | 0.07292 | 0.23438 | 1.58542 | NA |
|  | SLC2A6 | 0.01184 | 0.07392 | 0.03587 | 0.33919 | NA |
|  | LIF | 0.01251 | 0.07703 | 0.16888 | 0.79033 | NA |
|  | SOCS3 | 0.01269 | 0.07717 | 0.01914 | 0.98943 | NA |
|  | MLKL | 0.01288 | 0.07728 | 0.05028 | 0.69680 | NA |
|  | LOC132328383 | 0.01321 | 0.07826 | 0.12523 | 0.78551 | NA |
|  | TMC5 | 0.01373 | 0.08032 | 0.16570 | 1.36104 | NA |
|  | LOC132335646 | 0.01406 | 0.08120 | 0.08717 | 1.47927 | NA |
|  | PNP | 0.01585 | 0.08720 | 0.01243 | 0.69558 | NA |
|  | LOC132324463 | 0.01548 | 0.08720 | 0.09485 | 1.02348 | NA |
|  | LOC132325633 | 0.01571 | 0.08720 | 0.07163 | 0.32236 | NA |
|  | CLDN10 | 0.01666 | 0.08845 | 0.08385 | 0.92728 | NA |
|  | FSTL3 | 0.01660 | 0.08845 | 0.07646 | 1.04491 | NA |
|  | LIPI | 0.01795 | 0.09422 | 0.08185 | 1.64509 | NA |
|  | PIGR | 0.01922 | 0.09977 | 0.21244 | 5.28290 | NA |
|  | LOC132335743 | 0.01967 | 0.09981 | 0.54102 | 5.05700 | NA |
|  | LOC132329926 | 0.02009 | 0.09981 | 0.33647 | 3.48762 | NA |
|  | CD44 | 0.01994 | 0.09981 | 0.02707 | 0.13078 | NA |
|  | GPX3 | 0.01987 | 0.09981 | 0.09418 | 0.59345 | NA |
| **int+high > sham+high** | MYL2 | <0.00001 | 0.00043 | 0.02013 | NA | 0.67996 |
|  | C1QA | <0.00001 | 0.00843 | 0.05760 | NA | 1.16854 |
| **sham+high > low+high** | LOC132340237 | 0.00324 | 0.03989 | 0.60477 | 0.10546 | NA |
|  | LOC132332995 | 0.00503 | 0.04757 | 1.98791 | 0.01777 | NA |
|  | LOC132325636 | 0.00660 | 0.05055 | 0.74022 | 0.09170 | NA |
|  | LOC132332991 | 0.00690 | 0.05055 | 1.72769 | 0.43832 | NA |
|  | LOC132333048 | 0.01566 | 0.08720 | 1.01657 | 0.25012 | NA |
|  | LOC132332733 | 0.01640 | 0.08845 | 2.13296 | 0.46186 | NA |
| **sham+high > int+high** | LOC132340237 | 0.00324 | 0.03989 | 0.60477 | NA | 0.10546 |
|  | LOC132332995 | 0.00503 | 0.04757 | 1.98791 | NA | 0.01777 |

**Table S6**. Significantly differentially variable genes for MG infecting birds that received a secondary MG dose, where comparisons are between the sham+high treatment group versus the low+high or intermediate+high treatment groups. Loci are organized by ascending p-value/adjusted p-value and are listed if they had an adjusted p-value of <0.1. Data is grouped according to whether the variance was greater in the sham or prior exposed treatment (“Treatment Comparison”). Variance per gene per treatment group are reported unless a group was not involved in the statistical comparison (“na”). Unannotated loci (“HFMG94VAA”) have their predicted protein listed (“Product”).

| **Treatment Comparison** | **Locus** | **P-value** | **Adjusted P-value** | **Sham+High Variance** | **Low+High Variance** | **Int+High Variance** | **Product** |
| --- | --- | --- | --- | --- | --- | --- | --- |
| low+high > sham+high | lspA | >0.0001 | 0.0003 | 0.0371 | 1.4624 | NA |  |
|  | cdsA | >0.0001 | 0.0003 | 0.0476 | 1.9771 | NA |  |
|  | HFMG94VAA_RS02070 | >0.0001 | 0.0010 | 0.1878 | 2.7380 | NA | ABC transporter ATP-binding protein |
|  | ispF | >0.0001 | 0.0010 | 0.0713 | 1.4781 | NA |  |
|  | HFMG94VAA_RS02715 | >0.0001 | 0.0010 | 0.1085 | 3.0098 | NA | hypothetical protein |
|  | HFMG94VAA_RS00625 | >0.0001 | 0.0010 | 0.0930 | 1.4087 | NA | MPN157 family protein |
|  | rpoC | >0.0001 | 0.0010 | 0.0500 | 2.0005 | NA |  |
|  | HFMG94VAA_RS02210 | >0.0001 | 0.0012 | 0.0531 | 1.2887 | NA | hypothetical protein |
|  | ach1 | >0.0001 | 0.0012 | 0.0648 | 0.7845 | NA |  |
|  | HFMG94VAA_RS02300 | >0.0001 | 0.0012 | 0.0822 | 1.9023 | NA | ribosome biogenesis GTPase YqeH |
|  | folD | 0.0001 | 0.0027 | 0.1477 | 2.2649 | NA |  |
|  | HFMG94VAA_RS03095 | 0.0001 | 0.0027 | 0.0627 | 2.2184 | NA | DNA polymerase III subunit delta |
|  | acpS | 0.0001 | 0.0031 | 0.0661 | 1.0526 | NA |  |
|  | HFMG94VAA_RS03700 | 0.0001 | 0.0036 | 0.1019 | 1.8530 | NA | IS256 family transposase |
|  | HFMG94VAA_RS03065 | 0.0001 | 0.0036 | 0.2289 | 2.3771 | NA | ECF transporter S component |
|  | cysS | 0.0001 | 0.0036 | 0.1096 | 0.8179 | NA |  |
|  | smtA | 0.0002 | 0.0036 | 0.0950 | 1.2665 | NA |  |
|  | HFMG94VAA_RS02100 | 0.0002 | 0.0036 | 0.1119 | 1.3664 | NA | TIGR00282 family metallophosphoesterase |
|  | HFMG94VAA_RS01065 | 0.0002 | 0.0037 | 0.3649 | 3.4056 | NA | DNA-binding protein |
|  | smpB | 0.0002 | 0.0038 | 0.0405 | 1.2975 | NA |  |
|  | dX | 0.0003 | 0.0052 | 0.2694 | 3.1041 | NA |  |
|  | cdd | 0.0003 | 0.0052 | 0.0701 | 1.1944 | NA |  |
|  | rpsT | 0.0003 | 0.0054 | 0.1159 | 1.2510 | NA |  |
|  | prsA | 0.0003 | 0.0058 | 0.2119 | 1.8258 | NA |  |
|  | gapd | 0.0003 | 0.0058 | 0.0708 | 0.6995 | NA |  |
|  | HFMG94VAA_RS02775 | 0.0004 | 0.0059 | 0.1650 | 1.1258 | NA | HIT family protein |
|  | HFMG94VAA_RS02245 | 0.0005 | 0.0078 | 0.2466 | 2.8150 | NA | hypothetical protein |
|  | cmk | 0.0005 | 0.0081 | 0.2134 | 2.3777 | NA |  |
|  | dxr | 0.0006 | 0.0082 | 0.1418 | 1.2935 | NA |  |
|  | hrcA | 0.0009 | 0.0126 | 0.0909 | 2.1609 | NA |  |
|  | udk | 0.0009 | 0.0126 | 0.1202 | 1.3277 | NA |  |
|  | scpB | 0.0012 | 0.0153 | 0.2087 | 1.1806 | NA |  |
|  | HFMG94VAA_RS03485 | 0.0013 | 0.0156 | 0.0897 | 0.7785 | NA | serine/threonine protein kinase |
|  | ftsZ | 0.0016 | 0.0186 | 0.0179 | 0.3709 | NA |  |
|  | spxA | 0.0018 | 0.0202 | 0.1196 | 1.2647 | NA |  |
|  | HFMG94VAA_RS03635 | 0.0019 | 0.0207 | 0.3769 | 2.5600 | NA | hypothetical protein |
|  | HFMG94VAA_RS01490 | 0.0019 | 0.0207 | 0.2910 | 1.5744 | NA | sugar ABC transporter permease |
|  | metK | 0.0021 | 0.0214 | 0.0777 | 1.2483 | NA |  |
|  | HFMG94VAA_RS02425 | 0.0021 | 0.0214 | 0.2129 | 1.4561 | NA | aldo/keto reductase |
|  | HFMG94VAA_RS00785 | 0.0021 | 0.0214 | 0.2223 | 1.6724 | NA | DUF1951 domain-containing protein |
|  | HFMG94VAA_RS03130 | 0.0022 | 0.0214 | 0.3318 | 2.8608 | NA | ribonuclease M5 |
|  | HFMG94VAA_RS01205 | 0.0022 | 0.0214 | 0.1044 | 0.7902 | NA | FIVAR domain-containing protein |
|  | ksgA | 0.0023 | 0.0214 | 0.2972 | 1.7903 | NA |  |
|  | HFMG94VAA_RS00505 | 0.0023 | 0.0216 | 0.1414 | 1.0734 | NA | hypothetical protein |
|  | HFMG94VAA_RS02725 | 0.0025 | 0.0220 | 0.2375 | 1.3124 | NA | TrmH family RNA methyltransferase |
|  | atpH | 0.0025 | 0.0220 | 0.0559 | 0.6488 | NA |  |
|  | HFMG94VAA_RS03695 | 0.0026 | 0.0220 | 0.2792 | 2.0040 | NA | ATP-binding protein |
|  | lon | 0.0028 | 0.0233 | 0.0720 | 0.4585 | NA |  |
|  | trkG | 0.0035 | 0.0291 | 0.1303 | 0.8528 | NA |  |
|  | degV | 0.0040 | 0.0318 | 0.2006 | 1.0518 | NA |  |
|  | ispE | 0.0040 | 0.0318 | 0.3184 | 1.6099 | NA |  |
|  | HFMG94VAA_RS03675 | 0.0044 | 0.0346 | 0.2859 | 2.1569 | NA | 23S rRNA (guanosine(2251)-2'-O)-methyltransferase RlmB |
|  | rbgA | 0.0045 | 0.0346 | 0.1631 | 1.5551 | NA |  |
|  | HFMG94VAA_RS02085 | 0.0047 | 0.0353 | 0.2067 | 1.2905 | NA | IS256 family transposase |
|  | trpS | 0.0052 | 0.0385 | 0.2243 | 2.8587 | NA |  |
|  | HFMG94VAA_RS01470 | 0.0057 | 0.0416 | 0.2710 | 1.7157 | NA | hypothetical protein |
|  | HFMG94VAA_RS03995 | 0.0060 | 0.0418 | 0.2193 | 1.7207 | NA | IS1634 family transposase |
|  | plsX | 0.0060 | 0.0418 | 0.1949 | 1.2930 | NA |  |
|  | msrB | 0.0063 | 0.0418 | 0.2284 | 1.1779 | NA |  |
|  | HFMG94VAA_RS01440 | 0.0063 | 0.0418 | 0.2935 | 2.0172 | NA | hypothetical protein |
|  | rpmI | 0.0064 | 0.0418 | 0.1401 | 2.1423 | NA |  |
|  | mscL | 0.0064 | 0.0418 | 0.1286 | 2.2643 | NA |  |
|  | tmk | 0.0065 | 0.0418 | 0.6271 | 6.5971 | NA |  |
|  | exo | 0.0069 | 0.0439 | 0.1726 | 1.0108 | NA |  |
|  | rluA_1 | 0.0076 | 0.0479 | 0.0665 | 0.7853 | NA |  |
|  | cspR | 0.0080 | 0.0496 | 0.1790 | 0.9259 | NA |  |
|  | HFMG94VAA_RS00690 | 0.0082 | 0.0499 | 0.2753 | 1.6453 | NA | YhcH/YjgK/YiaL family protein |
|  | HFMG94VAA_RS04210 | 0.0084 | 0.0502 | 0.2785 | 2.3936 | NA | hypothetical protein |
|  | HFMG94VAA_RS00695 | 0.0089 | 0.0527 | 0.0383 | 1.5060 | NA | hypothetical protein |
|  | HFMG94VAA_RS03670 | 0.0090 | 0.0527 | 0.2115 | 1.1918 | NA | hypothetical protein |
|  | rplE | 0.0092 | 0.0529 | 0.0563 | 0.7024 | NA |  |
|  | HFMG94VAA_RS00410 | 0.0094 | 0.0535 | 0.9029 | 4.2236 | NA | hypothetical protein |
|  | HFMG94VAA_RS02785 | 0.0099 | 0.0550 | 0.1281 | 3.7040 | NA | transcription antitermination factor NusB |
|  | HFMG94VAA_RS01075 | 0.0100 | 0.0550 | 0.1937 | 1.5969 | NA | DUF3217 domain-containing protein |
|  | hemN | 0.0106 | 0.0564 | 0.4289 | 1.6090 | NA |  |
|  | pstS | 0.0106 | 0.0564 | 0.3783 | 3.4235 | NA |  |
|  | HFMG94VAA_RS03030 | 0.0106 | 0.0564 | 0.8145 | 3.5927 | NA | MFS transporter |
|  | HFMG94VAA_RS00495 | 0.0108 | 0.0569 | 0.1436 | 0.7726 | NA | hypothetical protein |
|  | rnhC | 0.0117 | 0.0604 | 0.2530 | 1.3447 | NA |  |
|  | HFMG94VAA_RS03520 | 0.0118 | 0.0604 | 0.1921 | 1.6754 | NA | AAA family ATPase |
|  | rpsB | 0.0122 | 0.0611 | 0.0533 | 0.2722 | NA |  |
|  | ffh | 0.0124 | 0.0611 | 0.1888 | 0.8257 | NA |  |
|  | HFMG94VAA_RS01765 | 0.0125 | 0.0611 | 0.2839 | 1.4789 | NA | DUF5454 family protein |
|  | glpF | 0.0125 | 0.0611 | 0.0872 | 1.1124 | NA |  |
|  | ribF/trmU | 0.0137 | 0.0659 | 0.2988 | 1.3739 | NA |  |
|  | cas2 | 0.0141 | 0.0669 | 0.2993 | 2.3053 | NA |  |
|  | HFMG94VAA_RS02440 | 0.0142 | 0.0669 | 0.6075 | 2.1449 | NA | hypothetical protein |
|  | HFMG94VAA_RS04190 | 0.0153 | 0.0713 | 0.1013 | 1.1299 | NA | hypothetical protein |
|  | HFMG94VAA_RS02905 | 0.0157 | 0.0724 | 0.1408 | 1.0115 | NA | hypothetical protein |
|  | HFMG94VAA_RS02680 | 0.0160 | 0.0727 | 0.5066 | 2.6343 | NA | hemolysin family protein |
|  | truB | 0.0164 | 0.0737 | 0.3365 | 1.4081 | NA |  |
|  | HFMG94VAA_RS03075 | 0.0168 | 0.0746 | 0.4008 | 2.6070 | NA | type I glyceraldehyde-3-phosphate dehydrogenase |
|  | gmk | 0.0176 | 0.0775 | 0.1957 | 0.9071 | NA |  |
|  | rpsU | 0.0182 | 0.0787 | 0.0804 | 0.8224 | NA |  |
|  | potE | 0.0182 | 0.0787 | 0.1048 | 1.3523 | NA |  |
|  | HFMG94VAA_RS04140 | 0.0217 | 0.0924 | 0.7444 | 2.9932 | NA | hypothetical protein |
|  | HFMG94VAA_RS00480 | 0.0219 | 0.0924 | 0.0812 | 0.3804 | NA | site-2 protease family protein |
|  | HFMG94VAA_RS03710 | 0.0224 | 0.0924 | 0.3017 | 1.8685 | NA | hypothetical protein |
|  | HFMG94VAA_RS04415 | 0.0225 | 0.0924 | 0.1652 | 0.8145 | NA | IS256 family transposase |
|  | HFMG94VAA_RS00120 | 0.0226 | 0.0924 | 0.7084 | 1.9436 | NA | hypothetical protein |
|  | HFMG94VAA_RS02685 | 0.0229 | 0.0924 | 0.6321 | 2.7527 | NA | deoxynucleoside kinase |
|  | HFMG94VAA_RS02335 | 0.0230 | 0.0924 | 0.1994 | 1.7016 | NA | hypothetical protein |
|  | HFMG94VAA_RS02975 | 0.0243 | 0.0967 | 0.3314 | 1.1860 | NA | potassium channel family protein |
| int+high > sham+high | HFMG94VAA_RS02340 | >0.0001 | 0.0004 | 0.0449 | NA | 2.3636 | FIVAR domain-containing protein |
|  | HFMG94VAA_RS03585 | 0.0001 | 0.0231 | 0.0706 | NA | 1.4404 | Ig-specific serine endopeptidase MIP |
|  | ptsG_1 | 0.0002 | 0.0321 | 0.0174 | NA | 2.6463 |  |
| sham+high > low+high | ugpC | 0.0011 | 0.0136 | 1.0407 | NA | 0.0548 |  |

**Table S7**. Enriched GO terms for significantly differentally variable (SDV) loci of MG infecting house finch conjunctiva. The gene list was made based on SDV loci identified between house finches that received a secondary dose of MG but were either naïve (sham+high) or had prior experience with the pathogen (low+high). GO terms are organized by ascending p-value/adjusted p-value. Terms are listed if they were enriched, regardless of significance. GO term categories were “MF” (Molecular Function), “CC” (Cellular Component), and “BP” (Biological Process). Gene ratio is the number of genes of interest that belong to a GO term out of all genes of interest (i.e., gene list). Background ratio is the number of genes per term out of all possible genes.

| GO Term | Description | Category | Gene Ratio | Background Ratio | P-value | Adjusted P-value |
| --- | --- | --- | --- | --- | --- | --- |
| GO:0046872 | metal ion binding | MF | 7/55 | 28/377 | 0.0946 | 0. 8946 |
| GO:0006508 | proteolysis | BP | 3/55 | 15/377 | 0.3780 | 0. 8946 |
| GO:0005737 | cytoplasm | CC | 12/55 | 75/377 | 0.4095 | 0. 8946 |
| GO:0005524 | ATP binding | MF | 9/55 | 56/377 | 0.4329 | 0. 8946 |
| GO:0005840 | ribosome | CC | 3/55 | 17/377 | 0.4629 | 0. 8946 |
| GO:1990904 | ribonucleoprotein complex | CC | 3/55 | 18/377 | 0.5036 | 0. 8946 |
| GO:0005525 | GTP binding | MF | 2/55 | 12/377 | 0.5436 | 0. 8946 |
| GO:0005829 | cytosol | CC | 8/55 | 58/377 | 0.6403 | 0. 8946 |
| GO:0016020 | membrane | CC | 2/55 | 15/377 | 0.6710 | 0. 8946 |
| GO:0016887 | ATP hydrolysis activity | MF | 4/55 | 32/377 | 0.7177 | 0. 8946 |
| GO:0008270 | zinc ion binding | MF | 2/55 | 17/377 | 0.7393 | 0. 8946 |
| GO:0005886 | plasma membrane or bacterial inner membrane | CC | 2/55 | 18/377 | 0.7688 | 0. 8946 |
| GO:0019843 | rRNA binding | MF | 3/55 | 30/377 | 0.8452 | 0. 8946 |
| GO:0006412 | translation | BP | 5/55 | 48/377 | 0.8658 | 0. 8946 |
| GO:0003735 | structural constituent of ribosome | MF | 1/55 | 14/377 | 0.8946 | 0. 8946 |
| GO:0022625 | cytosolic large ribosomal subunit | CC | 1/55 | 14/377 | 0.8946 | 0.8946 |

quantification of *Mycoplasma gallisepticum* genome load in conjunctival samples of

experimentally infected house finches (*Caprodacus mexicanus*) using real-time polymerase chain reaction. *Avian Pathology*, 37(4), 385-391.
